# Host and environmental factors differentially affect patterns of diversity in specialist and generalist parasites

**DOI:** 10.64898/2026.06.10.731414

**Authors:** Deesha Jeppu, Tanvi Kadakol, Nivedita Naveen, Guha Dharmarajan

## Abstract

**Aim:** Elucidating the mechanisms shaping parasite diversity patterns is critical because parasites encompass about 40% of known species, and are crucial for ecosystem structure and function. In free-living species, diversity patterns in the Anthropocene are shaped by niche-breadth because specialists (narrow niche-breadth taxa) are more sensitive to environmental disturbance compared to generalists (broad niche-breadth taxa). Like free-living species, parasites too can be categorized as specialists or generalists according to their niche-breadth (i.e., diversity of hosts they can infect). However, unlike free-living species, the effects of niche-breadth on parasite diversity patterns remain unclear. Here, we used haemosporidian parasites as a model system to identify factors affecting parasite diversity patterns, and test if these patterns differ between specialist (*Haemoproteus*) and generalist (*Plasmodium*) parasites.

**Location:** Southern India

**Taxon:** *Haemoproteus* spp. and *Plasmodium* spp. (Haemosporida)

**Methods:** Blood samples from wild birds were screened using molecular tools to identify haemosporidian parasite lineages. Statistical analyses, including random forest models and generalized dissimilarity models, were utilized to evaluate how environmental and host factors drive spatial patterns of parasite α and β diversity.

**Results:** Our results reveal that phylogenetic diversity is primarily shaped by host-related variables in the specialist parasites, but by numerous host- and environment-related factors in the generalists. In keeping with ecological theory, the specialist parasites showed higher α diversity and lower evenness compared to the generalists. Additionally, while β diversity of the specialist parasites was primarily driven by spatial differences in richness (e.g., taxon nestedness) rather than replacement (e.g., taxon turnover), the opposite pattern was found in the generalist.

**Main conclusion:** The differential patterns and drivers of diversity in specialist vs. generalist parasites demonstrates why specialists parasites are good indicators of ecosystem health and elucidates the mechanism by which anthropogenic disturbance increases the risk of emerging infectious diseases which are primarily caused by generalist parasites.

## 1 Introduction

Large-scale and rapid human-mediated modifications of natural environments is driving the catastrophic loss of species diversity in the Anthropocene (Ceballos et al., 2015; Turvey & Crees, 2019; Pyron & Pennell, 2022). However, extinction risk is not homogenous across species, with sensitivity to disturbance being greater in specialist taxa with narrow-niches compared to generalists with broad-niches (Kotiaho et al., 2005; Boyles & Storm, 2007; Devictor et al., 2007; Devictor et al., 2008; Liang et al., 2019; Cloyed et al., 2021; Moore et al., 2023).

Consequently, generalists tend to dominate communities in human-altered landscapes leading to the biotic homogenization of natural communities (McKinney & Lockwood, 1999; Olden et al., 2004; Devictor et al., 2007; Clavel et al., 2011). Biotic homogenization negatively impacts natural ecosystems (Olden et al., 2004; Clavel et al., 2011; Wang et al., 2021) and human populations (Olden et al., 2005). Critically, biotic homogenization can increase the risk of disease emergence in human-dominated landscapes (Gibb et al., 2020; Ostfeld & Keesing, 2020; Keesing & Ostfeld, 2021; Pei et al., 2025)

Emerging infectious diseases (EIDs) remain one of the greatest challenges of our time (Gupta et al., 2020; Dharmarajan et al., 2022). While parasites have been long recognized as causative agents of disease, they also play important roles in ecosystem structure and function (Hatcher et al., 2012b; Gupta et al., 2020). Parasites are good indicators of ecosystem health because of their diversity (Poulin & Morand, 2000; Dobson et al., 2008; Carlson et al., 2020), sensitivity to disturbance (Rezende et al., 2007; Dunn et al., 2009; Thompson et al., 2018) and myriad roles in maintaining ecosystem health (Marcogliese, 2005; Hudson et al., 2006; Betts et al., 2018; Frainer et al., 2018; Mougi, 2022; Brian, 2023). Despite their ecological importance parasites remain relatively understudied compared to free-living species (Gupta et al., 2020; Lymbery & Smit, 2023). Consequently, the mechanisms driving the structure of parasite communities remain unclear.

As in free-living species, a critical factor that can shape parasite community structure is niche-breadth, which in the case of parasites is related to the diversity of hosts they can infect. Based on niche-breadth, parasites can be categorized into specialist taxa that infect a few closely related host species and generalist taxa that can infect a wide range of hosts (Cooper et al., 2012; Gupta et al., 2020; Dharmarajan et al., 2021; Manzoli et al., 2021). Specialist parasites, show strong co-evolutionary dynamics with their hosts (Hudson et al., 2006) and increase host genetic diversity through negative frequency-dependent selection (i.e., Red Queen dynamics; Van Valen, 1973; Betts et al., 2018; Ebert, 2025). Such negative-frequency dependent selection also increases host species diversity by reducing local abundance of competitively dominant species (i.e., a Jenzen-Connell effect; Hudson et al., 2006; Terborgh, 2020). By directly driving host diversity specialist parasites help shape host community structure and promote ecosystem health (Fenton & Brockhurst, 2008). Specialist parasite are especially good indicators of ecosystem health because they are sensitive to disturbance (Lafferty, 2012; Dharmarajan et al., 2021) and can suffer co-extinction with their hosts (Dunn et al., 2009; Thompson et al., 2018; Carlson et al., 2020).

While habitat disturbance can negatively impact specialist parasites, they favor generalist parasites (Dharmarajan et al., 2021). The increased prevalence of generalist parasites has important implications for public health, because their broad niche-breadth allows such parasites to infect novel hosts, thus increasing the risk of EIDs (Timms & Read, 1999; Hatcher et al., 2012a; Farrell et al., 2013; Johnson et al., 2015; Wells & Clark, 2019; Dharmarajan et al., 2021). Additionally, generalist parasites are important from a conservation perspective because competent reservoir species (i.e., taxa that harbor the parasites but don’t suffer negative fitness consequences) can facilitate the maintenance of high prevalence of the parasite in the community which can lead to extirpation of susceptible species (Hudson et al., 2006; Gupta et al., 2020).

Since generalist parasites dominate host communities inhabiting disturbed habitats, parasite communities may undergo biotic homogenization, like free-living ones. The biotic homogenization of parasite communities is a cause for concern because the loss of specialist parasites negatively impacts ecosystem health while the increased prevalence of generalist parasites increases the risk of EIDs.

While several studies have focused on the drivers of parasite diversity (Morand & Poulin, 2000; Thieltges et al., 2011; Kamiya et al., 2014; Martins et al., 2021; McNew et al., 2021; Brian & Aldridge, 2023; Dutra, 2023; Brian & Aldridge, 2024), it remains unclear if these drivers differ between specialist vs. generalist parasites. In this study, we address this gap using, as a model system, avian haemosporidian parasites infecting birds inhabiting the sky-islands of the Western Ghats, Southern India. Avian haemosporidian parasites are transmitted by dipteran vectors to numerous bird species globally, reducing their lifespan and fitness (Atkinson & Samuel, 2010; Asghar et al., 2015; Ilgūnas et al., 2016; Schoepf et al., 2022). Avian haemosporidians are an ideal model system to address the objectives of our study because of the high diversity of parasite lineages and host species they infect (Beadell et al., 2009; Pigeault et al., 2015; Gupta et al., 2019). Additionally, the two major genera (*Haemoproteus* and *Plasmodium*) show distinct differences in their host breadth, with lineages of *Haemoproteus* being greater host specialists compared to *Plasmodium* in our study system (Gupta et al., 2019), as well as a large number of other systems globally (Waldenstrom et al., 2002; Beadell et al., 2004; Fallon et al., 2005; Beadell et al., 2009; Dimitrov et al., 2010; Okanga et al., 2014; Musa et al., 2019; Doussang et al., 2021; Ndlovu et al., 2024; Cebrian-Camison et al., 2025), though some exceptions exist (e.g., bird communities sampled in some parts of the Neotropics; see Moens & Perez-Tris, 2016; Moens et al., 2016).

In this study we identify the main drivers of α and β diversity (Whittaker, 1960; Arellano & Halffter, 2022) in the host (bird) and parasite (Haemosporidian) communities, and specifically focus on understanding how these drivers differ between specialist and generalist parasites (i.e., *Haemoproteus* and *Plasmodium*, respectively). The shola sky-islands of the southern Western Ghats are a biodiversity hotspot characterized by recent land-use change driven by anthropogenic factors, such as urbanization, sylviculture and agriculture (Robin & Nandini, 2012). The bird communities in this region are characterized by high levels of endemism, as well as a combination of both range-restricted ancient and young lineages (Goyal et al., 2025).

Consequently, we expected the bird communities would be characterized by high levels of α diversity but low levels of α evenness. Given the high levels of human-mediated disturbance in conjunction with natural heterogeneity (e.g., elevation gradients) we also expected that the bird communities would show high levels of β diversity. However, we expected that species loss would be non-random with the preferential loss of specialist species (e.g., high-elevation and/or shola habitat specialists) compared to generalist species. Consequently, we expected bird communities would show a strongly nested community structure, and that overall β diversity would primarily be driven by differences in species richness (i.e., due to nestedness) rather than species replacement (i.e., spatial turnover) between communities (Cardoso et al., 2014).

Ecological theory predicts that biogeographical differences between specialist vs. generalist taxa will be critically impacted by the differential extent of their reliance on a specific resources due to differences in their niche-breadths (Carscadden et al., 2020). Based on niche theory, in the case of parasites, we predicted that specialist parasites would be more strongly impacted by community structure of their hosts compared to generalist parasites because the distribution of the former would be highly correlated with the distribution of the specific resources (i.e., host species) they rely on (Dunn et al., 2009; Thompson et al., 2018; Carlson et al., 2020).

Consequently, we expected that the specialist parasite (*Haemoproteus*) would show high levels α diversity (i.e., a large number of lineages each infecting only a small fraction of host species) but low levels of evenness (i.e., high variability in lineage prevalence across sites) compared to the generalist parasite (*Plasmodium*). We also expected that, like the host communities, *Haemoproteus* β diversity would be dominated by changes in parasite lineage richness (i.e., due to taxon nestedness) while that of the generalist *Plasmodium* would be dominated by parasite lineage replacement (i.e., due to taxon turnover). Finally, we expect that α and β diversity of specialist *Haemoproteus* would be driven primarily by host-related factors (e.g., host phylogenetic and functional diversity), while that of generalist *Plasmodium* would be driven by both host and environmental factors (e.g., temperature and rainfall).

## 2 Methods and Methods

### 2.1 Databases

This study used ecological and environmental data that were derived from previous studies and/or public databases.

#### 2.1.1 Ecological data

Bird and parasite genetic data for this study were obtained from DNA extracted from blood samples (N = 1172) collected from 28 species of birds trapped from the Western Ghats, Southern India. The fieldwork related to sample collection was carried out at 42 sites distributed across four geographical regions spanning the southern 600 km of the Western Ghats (Table S1). Adult birds were mist-netted during the pre-monsoon, pre-breeding season (January–May) between 2011 and 2013 following the methods reported earlier (Robin et al., 2010). Blood samples (50– 100 μl) were drawn from the ulnar vein using heparinized capillary tubes and stored in Queen’s lysis buffer at room temperature during fieldwork, then transferred to –20°C in the laboratory (Seutin et al., 1991). The bird phylogenetic tree were constructed from cytochrome b sequence data (1143bp) using MrBayes ver 3.1.2 (Ronquist & Huelsenbeck, 2003) as described in Robin et al. (2015). The bird functional diversity was derived from the Avonet database (Tobias et al., 2022). The parasite data were obtained from Gupta et al. (2019). Briefly, a nested PCR was used to amplify a 478-bp mitochondrial cytochrome b fragment of avian haemosporidian parasites from the bird blood samples (Hellgren et al., 2004). *Plasmodium relictum* genomic DNA served as a positive control. Amplicons were verified on agarose gel and parasite positive samples were bidirectionally sequenced using Sanger sequencing. Sequences were assembled in Geneious 9.1.5 (Kearse et al., 2012), and those with double peaks (mixed infections) were excluded from downstream analyses. Lineages were identified via GenBank and MalAvi databases (Bensch et al., 2009), with a ≥ 1 bp difference defining distinct lineages (Bensch et al., 2004). The parasite phylogenetic tree were constructed from cytochrome b sequence data using MrBayes ver 3.1.2 as described in Gupta et al. (2019).

#### 2.1.2 Environmental data

A key objective of this study was to examine how climate, landscape, and human disturbance affect patterns of parasite diversity, both directly and through their influence on host diversity (see details below). Climatic data was obtained in the form of 19 standard bioclimatic variables from http://chelsa-climate.org/bioclim/ (See Table S2 for details). The terrain metrics (i.e., elevation and slope) were extracted from a Digital Elevation Model (http://www.earthenv.org/DEM) using the R package RASTER. We also obtained data on water flow accumulation (https://www.hydrosheds.org/) and canopy height (http://lidarradar.jpl.nasa.gov). Landcover data was obtained from Roy et al. (2015) and we used these data to extract the proportions of 8 major habitat types (i.e., cropland, degraded habitat, forest, grassland, plantation, settlements, shrub/savannah and water). Anthropogenic disturbance was represented by distance from protected area boundaries (http://datasets.wri.org) and a human proximity index derived from Indian census data (http://www.ciesin.columbia.edu/data/india-census-grids/) following Alexander and Wint (2013). Details of environmental variables used are given in Table S2.

### 2.2 Statistical Methods

All statistical analyses were carried out using R ver. R version 4.4.1 (The R Foundation for Statistical Computing 2019), and all statistical tests and reported *P* values are two sided. A complete list of R packages used for analyses is given in Table S3.

#### 2.2.1 α Diversity Analyses

We used the α diversity and evenness functions in the R package BAT (Cardoso et al., 2014) to calculate the observed parasite phylogenetic α diversity and evenness for each site based on the observed parasite lineages identified in each site in conjunction with the parasite phylogenetic tree. In the case of avian hosts, we calculated both phylogenetic and functional diversity using the R package BAT. Phylogenetic diversity was based on the bird phylogenetic tree and functional diversity derived from Gower distances based on traits related to feeding strata, sociality, habitat use, and connectivity from the bird functional traits database (see section Databases above). Variable details are provided in Table S2. We tested if the mean α diversity and evenness across sites differed between *Plasmodium* and *Haemoproteus* using a t test as implemented in R. Finally, we tested the effects of biotic and environmental variables on the spatial patterns of parasite α diversity separately in *Plasmodium* and *Haemoproteus* using random forest models. Briefly, the random forests models were implemented in the R package RANGER (Wright & Ziegler, 2017). All RFMs were run using 100,000 trees and we optimized the model parameters (mTry, Min. node size, Split rule) using a 10-fold cross-validation procedure implemented in the trainControl function from the R package CARET (Kuhn, 2008). All RFMs also included the number of birds sampled in each trap site as case weights to control for potential sample size effects. We initially included all environmental variables for all RFMs, and also added host-related variables for the parasite-related RFMs. Thus, initially environmental factors (i.e., terrain, climate, habitat and disturbance) were included in the RFMs to predict phylogenetic and functional host diversity. In turn, parasite-related variables (parasite phylogenetic diversity) were assumed to be driven by direct effects of environmental variables and the indirect effects of these variables through their effects on host diversity. For each RFM, we first fit the full random forest model with all variables (Table S2), and estimated the variable importance values (Wright & Ziegler, 2017). We then used a forward step-wise selection starting from the most important variable, and adding variables in decreasing order of importance. At each step a variable was added if it’s correlation with variables included in the model until that step was ≤ 0.7, and if addition of the variable increased the coefficient of determination (R^2^) for the whole model. Model validation was done using three repetitions of a 10-fold cross validation procedure to maximize the R^2^ value of the whole model using 75% of the data to train the model and 25% to assess model prediction accuracy (Jacob et al., 2020). In order to calculate the relative importance of each of the four main variable groups (i.e., terrain, climate, habitat and ecology; Table S2) we also calculated a group variable importance measure. This measure was calculated as the weighted mean of the number of times variables in each group were used to split trees in the random forest (see https://stats.stackexchange.com/questions/92419). Finally, to facilitate interpretation of the patterns, we used fuzzy clustering across all raster cells using the function ‘fanny’ in the R package CLUSTER (Maechler ET AL., 2023) to identify three clusters based on the predicted α diversity. We tested the effectiveness of the clustering by evaluating if the identified clusters differed significantly in terms of α diversity using a Generalized Least Squares (GLS) model that controlled for spatial autocorrelation using the function ‘gls’ in the R package NLME (Pinheiro & Bates, 2000; Pinheiro et al., 2023). Further, we evaluated if the average values of the landscape variables in each cluster differed from each other using GLS models (see above).

#### 2.2.2 β Diversity Analyses

We used the beta function in the R package BAT (Cardoso et al., 2014) to calculate the observed parasite phylogenetic β diversity for each pair of sites based on the observed parasite lineages identified in each site in conjunction with the parasite phylogenetic tree. In the case of avian hosts, we calculated both phylogenetic and functional β diversity using the R package BAT. Phylogenetic diversity was based on the bird phylogenetic tree and functional diversity derived from Gower distances (see details above). We further partitioned the total β diversity (β_TOTAL_) into β_REPL_ (i.e., β diversity explained by replacement of species alone) and β_RICH_ (i.e., β diversity explained by richness differences alone) using the approaches described by Cardoso et al. (2014), as implemented in BAT. We tested if the mean pairwise β_REPL_ and β_RICH_ across sites differed between *Plasmodium* and *Haemoproteus* using the non-parametric U-test as implemented in R. Finally, we tested the effects of biotic and environmental variables on the spatial patterns of parasite α diversity separately in *Plasmodium* and *Haemoproteus* using generalized difference models (GDMs; Ferrier et al., 2007), as implemented via the GDM package in R (Fitzpatrick et al., 2021). All models used three i-spline basis functions per predictor, and model performance was evaluated through cross-validation, whereby 80% of sites were randomly selected to train the model and the remaining 20% used for validation; this process was repeated 10 times per predictor combination. Candidate predictors included geographic distance alongside the biotic and environmental variables used previously in the random forest models. As with the α diversity analysis, we first screened predictors based on individual explanatory power and inter-predictor correlations, then applied backward elimination to arrive at a parsimonious final model containing only statistically significant variables, confirmed via permutation testing (Fitzpatrick et al., 2021). In order to calculate the relative importance of each of the four main variable groups (i.e., terrain, climate, habitat and ecology; Table S2) we also calculated we also calculated a group variable importance measure. To calculate the measure, we calculated the relative change in deviance when all variables in a specific group were removed from the model compared to the full model. Final models of compositional dissimilarity were spatially projected using the final GDM model in conjunction with raster layers of the selected predictors. To visualize geographic patterns in community dissimilarity, we applied a principal components analysis (PCA) to the GDM-transformed predictor layers, assigned red, green, and blue color channels to the first three PCA axes, and combined these into a single map (Mokany et al., 2022). Finally, to facilitate interpretation of the patterns, we first calculated the predicted dissimilarity between each pair of raster cells. We then used this dissimilarity matrix to carry out a maximum of three fuzzy clusters using the ‘fanny’ function in the R package CLUSTER. (MAECHLER ET AL., 2023) based on the predicted α diversity. We tested the effectiveness of the clustering by evaluating if the dissimilarity between raster cells in the same cluster were significantly lower than that between raster cells in different clusters using a partial mantel test. The partial mantel test was used so as to control for the geographic distance between the raster cells, and was carried out using the R package VEGAN (Dixon, 2003). Finally, we evaluated if the average values of the landscape variables in each cluster differed from each other using GLS models that controlled for spatial autocorrelation (see details above).

## 3 Results

Our analysis included samples from a total of 1172 birds from 42 sites, ranging in elevation from 875 to 2256 m in elevation (Fig. S1; Table S1). In these samples, a total of 47 unique parasite lineages infecting birds in the study area were identified: 29 *Haemoproteus* lineages and 18 *Plasmodium* lineages.

### 3.1 α Diversity

We found that patterns of α diversity and evenness were in keeping with differences in parasite niche-breadth and niche theory expectations. Specifically, we found that across all sites, phylogenetic α diversity was significantly higher in the specialist (*Haemoproteus*) compared to the generalist (*Plasmodium*) parasite (Fig. 1a; t = 4.229; df = 58.958; *P* < 0.001). Alternatively, with phylogenetic α evenness, we found the opposite pattern, with *Haemoproteus* showing significantly lower levels of evenness compared to *Plasmodium* (Fig. 1b; t = -2.154; df = 31.815; *P* = 0.039). Spatial patterns related to α diversity also revealed clear differences between the two parasite taxa. With respect to *Haemoproteus* α diversity (α_H-PHYLO_) the RFM generally performed explaining about 77% of the variance amongst sites (Fig. 1c; Table S4; r^2^ ± SE = 0.774 ± 0.307). The final RFM retained the following variables: α_HST-PHYLO_, α_HST-FUNCT_ and PA_DIST_ (Table S5; Fig. S03). Thus, α_H-PHYLO_ was primarily affected by factors related to host ecology and disturbance, and these factors contributed to about 78% and 22% of the total variable importance, respectively (Fig. 1d). The predicted spatial patterns of α_H-PHYLO_ showed high levels of heterogeneity, with few isolated hot-spots of high diversity and large areas associated with low diversity (Fig. 1e). Fuzzy clustering based on patterns of α_H-PHYLO_ (Fig. 1e, inset maps) identified three clusters with α_H-PHYLO_ decreasing from cluster C1 to C3 (Fig. 1f; C1 > C2 > C3; χ^2^ = 4629.527; df = 2; *P* < 0.001). Additionally, raster cells sampled from the clusters differed with respect to key underlying drivers, including: α_HST-PHYLO_ (Fig. 1g; C1 > C2 > C3; χ^2^ = 491.727; df = 2; *P* < 0.001), α_HST-FUNCT_ (Fig. 1h; C1 > C2 > C3; χ^2^ = 323.854; df = 2; *P* < 0.001) and PA_DIST_ (Fig. 1i; C1 < C2 < C3; χ^2^ = 12.091; df = 2; *P* = 0.002).

**Figure 1.**
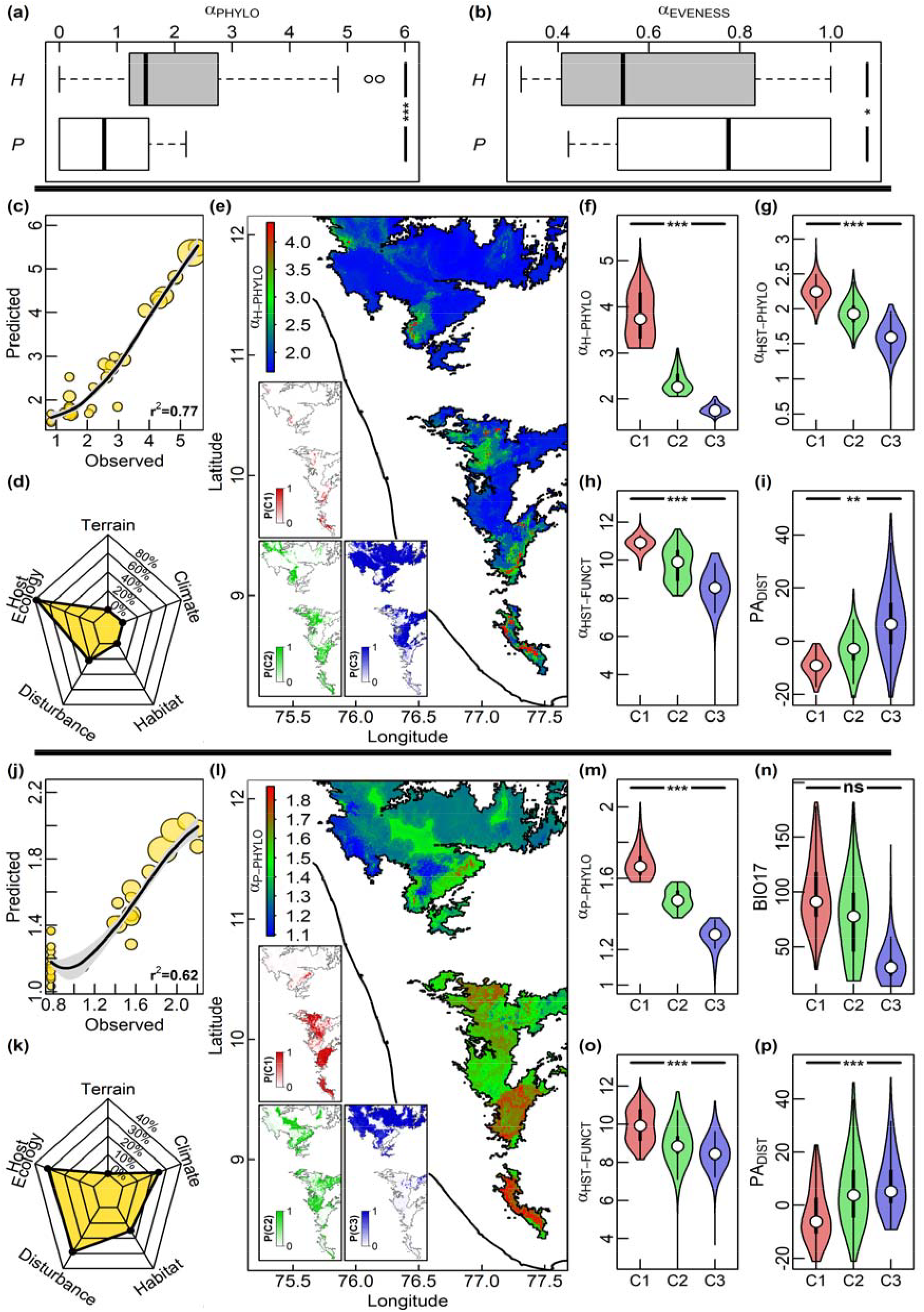
Avian haemosporidian parasite α diversity patterns in the Western Ghats, Southern India. (a) Across all sites, phylogenetic α diversity (α_PHYLO_) is higher in *Haemoproteus* (gray) vs. *Plasmodium*(white). (b) Alternatively, phylogenetic α evenness (α_EVENNESS_) is lower in *Haemoproteus* (gray)vs. *Plasmodium* (white). Spatial patterns of *Haemoproteus* phylogenetic α diversity (α_H-PHYLO_) based on the best fit Random Forests Model (RFM) analysis (c-e). (c) Observed vs. predicted scatter plot for α_H-PHYLO_, with size of points scaled relative to sample size in each site. (d) Relative importance of variables grouped into five major categories (terrain, climate-, habitat-, disturbance- and host ecology-related factors) in predicting α_H-PHYLO_. (e) Predicted spatial patterns of α_H-PHYLO_ across the Western Ghats study area. Inset maps show the geographical regions resulting from the Fuzzy C-Means (FCM) cluster analysis of α_H-PHYLO_ showing the probability of membership in the identified clusters C1 (red), C2 (green) and C3 (blue), respectively. For each cluster (C1-C3), the distribution of values are plotted as violin plots (f-i) for: (f) The response variable (α_H-PHYLO_), (g) host phylogenetic α diversity (α_HST-PHYLO_), (h) host functional α diversity (α_HST-FUNCT_), (i) distance to protected areas (PA_DIST_). Spatial patterns of *Plasmodium* phylogenetic α diversity (α_P-PHYLO_) based on the best fit RFM analysis (j-l). (j) Observed vs. predicted scatter plot for α_P-PHYLO_, with size of points scaled relative to sample size in each site. (k) Relative importance of variables grouped into five major categories (see Fig. 1d for details) in predicting α_P-PHYLO_. (l) Predicted spatial patterns of α_P-PHYLO_ across the Western Ghats study area, with inset maps showing results of FCM analysis (as in Fig. 1e). For each cluster (C1-C3), the distribution of values are plotted as violin plots (m-p) for: (m) The response variable (α_P-PHYLO_), (n) precipitation of driest quarter (BIO17), (o) host functional α diversity (α_HST-FUNCT_), (p) distance to protected areas (PA_DIST_). For box plots, the thick horizontal line and box represent the median and quartiles, respectively. Whiskers extend to 1.5 times the inter-quartile range and open circles represent outliers. For violin plots, the white dot and thick line represent the median and quartiles, respectively. The thin lines extend to 1.5 times the inter-quartile range. Significant differences between groups in box and violin plots are indicated by the following symbols: *** (*P* < 0.001), ** (*P* < 0.01), * (*P* < 0.05), ns (non-significant). See text for statistical test details.

With respect to *Plasmodium* α diversity (α_P-PHYLO_) the best-fit RFM for α_P-PHYLO_ explained about 62% of the variance amongst sites (Fig. 1j; Table S4; r^2^ ± SE = 0.622 ± 0.376). The final model retained the following variables: α_HST-FUNCT_, PA_DIST_, BIO17, BIO01, Degrad_PROP_ and BIO04 (Table S5; Fig. S04). Thus, α_P-PHYLO_ was affected by multiple types of factors, and factors related to ecology, disturbance, climate and habitat contributed to about 31%, 29%, 25% and 15% of the total variable importance, respectively (Fig. 1k). The predicted spatial patterns related to α_P-PHYLO_ showed relatively low levels of heterogeneity (Fig. 1l), compared to those associated with *Haemoproteus* (Fig. 1e). Fuzzy clustering based on patterns of α_P-PHYLO_ (Fig. 1l, inset maps) identified three clusters with α_P-PHYLO_ decreasing from cluster C1 to C3 (Fig. 1m; C1 > C2 > C3; χ^2^ = 752.065; df = 2; *P* < 0.001). Contrary to *Haemoproteus*, in the case of *Plasmodium* only a few independent variables differed significantly between the clusters. Specifically, raster cells sampled from the clusters did not differ with respect to four variables: BIO17 (Fig. 1n; χ^2^ = 4.645; df = 2; *P* = 0.098), BIO01 (χ^2^ = 1.511; df = 2; *P* = 0.47), BIO04 (χ^2^ = 1.007; df = 2; *P* = 0.605) and Degrad_PROP_ (χ^2^ = 3.122; df = 2; *P* = 0.21). Alternatively, we found that raster cells sampled from the clusters differed significantly with respect to two variables: α_HST-FUNCT_ (Fig. 1o; C1 > C2 > C3; χ^2^ = 105.817; df = 2; *P* < 0.001) and PA_DIST_ (Fig. 1p; C1 < C2 < C3; χ^2^ = 21.885; df = 2; *P* < 0.001). As expected, we see a strong concordance between parasite α diversity and host functional α diversity (α_HST-FUNCT_; Table S4-S5; Fig. S05a-g and S06) and/or phylogenetic α diversity (α_HST-PHYLO_; Table S4-S5; Fig. S05h-n and S07), though these signatures of concordance are stronger in the case of the specialist *Haemoproteus* vs. generalist *Plasmodium*.

### 3.2 β Diversity

Our analyses demonstrated that β diversity patterns, as in the case of α diversity, differed between *Haemoproteus* and *Plasmodium* lineages, and that these differences were in keeping with the theoretical expectations based on differences in parasite niche-breadth between these two parasite taxa. Specifically, with respect to β diversity, we found that across all sites, phylogenetic β diversity due to changes in richness was significantly higher in *Haemoproteus* vs. *Plasmodium* (Fig. 2a; W = 163272; *P* < 0.001). Alternatively, we found that across all sites, phylogenetic β diversity due to taxon replacement was significantly lower in *Haemoproteus* vs. *Plasmodium* (Fig. 2b; W = 91808; *P* < 0.001).

**Figure 2.**
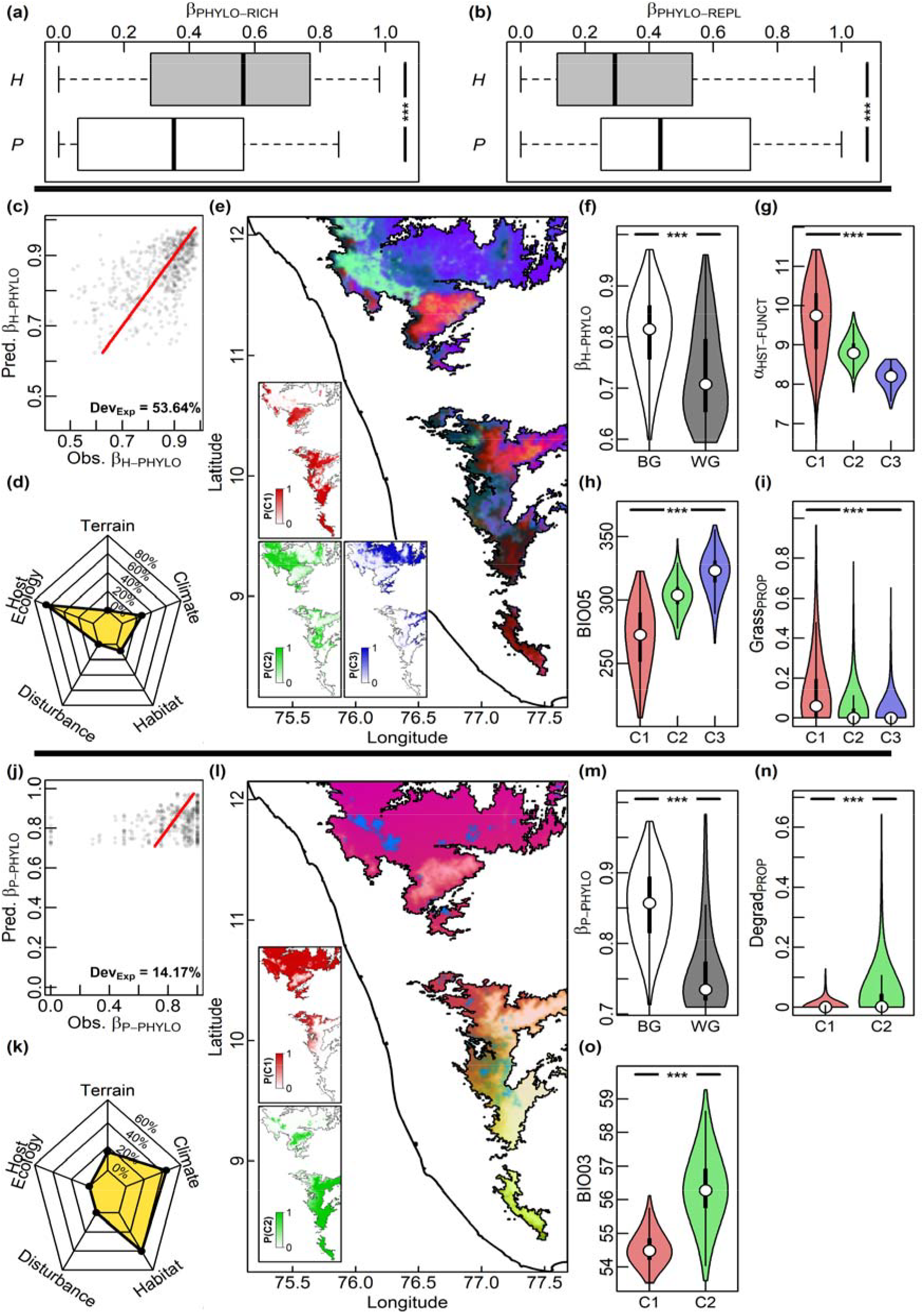
Avian haemosporidian parasite β diversity patterns in the Western Ghats, Southern India. (a) Across all sites, phylogenetic β diversity due to changes in richness (β_PHYLO-RICH_) is higher in *Haemoproteus* (gray) vs. *Plasmodium* (white). (b) Alternatively, phylogenetic β diversity due to taxon replacement (β_PHYLO-REPL_) is lower in *Haemoproteus* (gray) vs. *Plasmodium* (white). Spatial patterns of *Haemoproteus* phylogenetic total β diversity (β_H-PHYLO_) based on the best fit generalized difference model (GDM) analysis (c-e). (c) Observed dissimilarity as a function of GDM-predicted dissimilarity for β_H-PHYLO_. Each site-pair is represented as a point and the line of is equality provided. (d) Relative importance of variables grouped into five major categories (terrain, climate-, habitat-, disturbance- and host ecology-related factors) in predicting β_H-PHYLO_. (e) Predicted spatial patterns of β_H-PHYLO_ across the Western Ghats study area. Inset maps show the geographical regions resulting from the Fuzzy C-Means (FCM) cluster analysis of β_H-PHYLO_ showing the probability of membership in the identified clusters C1 (red), C2 (green) and C3 (blue), respectively. For each cluster (C1-C3), the distribution of values are plotted as violin plots (f-i) for: (f) The response variable (β_H-PHYLO_), (g) host functional α diversity (α_HST-FUNCT_), (h) max temperature of warmest month (BIO05), (i) proportion grassland (Grass_PROP_). Spatial patterns of *Plasmodium* phylogenetic total β diversity (β_P-PHYLO_) based on the best fit GDM analysis (j-l). (j) Observed dissimilarity as a function of GDM-predicted dissimilarity for β_P-PHYLO_. Each site-pair is represented as a point and the line of is equality provided. (k) Relative importance of variables grouped into five major categories (terrain, climate-, habitat-, disturbance- and host ecology-related factors) in predicting β_P-PHYLO_. (l) Predicted spatial patterns of β_P-PHYLO_ across the Western Ghats study area, with inset maps showing results of FCM analysis (as in Fig. 1e). For each cluster (C1-C2), the distribution of values are plotted as violin plots (m-p) for: (m) The response variable (β_P-PHYLO_), (n) proportion degraded habitat (Degrad_PROP_), (o) isothermality (BIO03). Box plot, violin plot and significance test details are as mentioned in Fig. 1. See text for statistical test details.

Generalized difference modelling (GDM) analyses also demonstrated that the two parasite taxa had differential spatial β diversity patterns. The best-fit generalized difference model (GDM) for β_H-PHYLO_ explained about 54% of the total deviance (Fig. 2c; Table S6; Model *P* value < 0.001), and retained the following variables: Grass_PROP_, Plantn_PROP_, BIO05 and α_HST-FUNCT_ (Fig. 2d; Table S7; Fig. S08). The predicted spatial patterns of *Haemoproteus* β diversity showed strong and relatively clear breaks that were independent of spatial proximity of the cells (Fig. 1e). As in the case of the α diversity analyses, we found that fuzzy clustering in the case of performed well with average β_H-PHYLO_ between groups being significantly greater than the average β_H-PHYLO_ within groups (Fig. 1f; Mantel test Spearman’s ρ = 0.384; *P* = 0.001). Additionally, we found that raster cells sampled from the clusters formed based on β_H-PHYLO_ differed with respect to key underlying drivers, including: α_HST-FUNCT_ (Fig. 1g; C1 > C2 > C3; χ^2^ = 459.869; df = 2; *P* < 0.001), BIO05 (Fig. 1h; C1 < C2 < C3; χ^2^ = 334.448; df = 2; *P* < 0.001), Grass_PROP_ (Fig. 1i; C1 > C2 > C3; χ^2^ = 71.562; df = 2; *P* < 0.001) and Plantn_PROP_ (C1 < C2; C1 > C3; χ^2^ = 65.476; df = 2; *P* < 0.001).

For β_P-PHYLO_, the best-fit GDM performed relatively poorly, and only explained about 14% of the total deviance (Fig. 2j; Table S6; Model *P* value < 0.001). The best-fit GDM retained (in order of importance) the following variables: Degrad_PROP_, BIO03 and geographic distance (Fig. 1k; Table S7; Fig. S09). The predicted β_P-PHYLO_ spatial patterns showed relatively weak patterns (Fig. 1l) compared to β_H-PHYLO_ (Fig. 1e), and unlike in the case of β_H-PHYLO_, these patterns were affected by the spatial proximity of the cells (Fig. 1k). Unlike the case of β_H-PHYLO_, fuzzy clustering only found evidence for two clusters in the case of β_P-PHYLO_. However, we found that the average β_P-PHYLO_ between groups was significantly greater than the average β_P-PHYLO_ within groups (Fig. 1m; Mantel test Spearman’s ρ = 0.509; *P* = 0.001), revealing that the clustering analysis performed well. Additionally, we found that raster cells sampled from the clusters formed based on β_P-PHYLO_ differed with respect to key underlying drivers, including: Degrad_PROP_ (Fig. 1n; C1 < C2; χ^2^ = 212.169; df = 1; *P* < 0.001) and BIO03 (Fig. 1o; C1 < C2; χ^2^ = 80.149; df = 1; *P* < 0.001). As expected, we see a strong concordance between parasite β diversity and host functional β diversity (β _HST-FUNCT_; Table S6-S7; Fig. S10a-g and S11) and/or phylogenetic β diversity (β _HST-PHYLO_; Table S6-S7; Fig. S10h-n and S12). However, as in the case of parasite α diversity, this concordance in the case of parasite β diversity was stronger in the case of the specialist *Haemoproteus* vs. generalist *Plasmodium*.

## 4 Discussion

In this study, we show that the drivers of specialist and generalist haemosporidian parasite diversity differ, presenting broad and significant implications for their roles in the ecosystem, parasite conservation, and the possible spread of EIDs. Specifically, with regard to α and β diversity, we find that host and environmental factors play differential roles of varying importance in predicting diversity and spatial patterns in specialist (*Haemoproteus*) vs. generalist (*Plasmodium*) parasites.

Our analysis demonstrated that phylogenetic α diversity was significantly higher in the specialist *Haemoproteus* compared to the generalist *Plasmodium*. Similar patterns of higher diversity amongst specialists vs. generalist taxa have been reported in free-living species, and a negative correlation between niche-breadth and diversity seems to be a general pattern observed across numerous taxa, independent of environmental characteristics or the diversity measure used (Granot & Belmaker, 2020). Niche-breadth is both a cause and a consequence of diversity, and that specialists are expected to show higher levels of diversity compared to generalists. For example, the classical theory of limiting similarity (Macarthur & Levins, 1967) predicts higher diversity in specialists vs. generalists because the narrow niche-breadth of the former allows for more efficient niche packing (MacArthur, 1965; Pigot et al., 2016; Pellissier et al., 2018; PaganiLJNúñez et al., 2019). There is also evidence that “diversity-begets-diversity” because increased species diversity and stronger biotic interactions can promote further diversification either via the creation of novel niches or through niche partitioning (Whittaker, 1972; Calcagno et al., 2017). Finally, in human-dominated landscapes, specialists will be expected to have especially low levels of diversity (compared to generalists) because specialists tend to have low tolerance to disturbance, and thus face increased vulnerability to extinction (McKinney & Lockwood, 1999).

Alternatively, with phylogenetic α evenness, we found the opposite pattern, with *Haemoproteus* showing significantly lower levels of evenness compared to *Plasmodium*. It is well-known that niche-breadth impacts the geographic distributions of species (Carscadden et al., 2020). In most ecological communities, concepts of rarity or commonness are inherently scale-dependent (Hartley & Kunin, 2003). For example, specialist taxa, which require specialized resources, tend to have narrow geographic distributions (Verberk et al., 2010; Slatyer et al., 2013) and high sensitivity to disturbance (Clavel et al., 2011) compared to generalist taxa. However, such specialists likely are competitively dominant under ideal conditions, and can thus reach very high abundances in their optimal habitats (Lesica et al., 2006). These differences in competitive ability and dispersal between specialists and generalists predicts an overall increase in evenness as disturbance increases (Freeman et al., 2018; Lorel et al., 2021), with specialist-dominated communities showing lower levels of evenness compared to generalist-dominated communities (Lôbo et al., 2011; Carrara et al., 2015; Ibarra & Martin, 2015).

Phylogenetic α diversity of *Haemoproteus* was primarily affected by factors related to host ecology and disturbance, while phylogenetic α diversity of *Plasmodium* was affected by a wider range of factors relating to ecology, disturbance, climate and habitat. Contrary to generalists, the α diversity of specialist parasites is more likely to be largely affected by host variables than environmental variables, as they have specialized mutualistic relationships due to coevolution with their hosts, and they can’t rapidly switch to a new host (Hellgren et al., 2008). This is not to say that ecology, disturbance, habitat and climatic variables do not impact specialist parasite α diversity, rather that such an impact would be indirect via their effects on host community structure, which explains why patterns of bird diversity are more congruent with *Haemoproteus* vs. *Plasmodium*. Specialist *Haemoproteus* and generalist *Plasmodium*, also differ strikingly with regard to the spatial heterogeneity in the magnitude of α diversity observed across our study area. Specifically, *Haemoproteus* shows high levels of spatial heterogeneity – with a few isolated hotspots of high diversity and large areas associated with low diversity, while *Plasmodium* α diversity shows low spatial heterogeneity. These results align with the general understanding of the relationship between niche-breadth and spatial patterns, where a positive correlation between niche-breadth and range size has been found across niche-breadth-related studies (Cardillo et al., 2018; Moulatlet et al., 2025). Critically, hotspots of hyper diversity of specialists should be protected, since any disturbance here could lead to a greater loss of diversity compared to sites with low diversity. Additionally, since spatial patterns for the specialist *Haemoproteus* match the diversity patterns of hosts, conserving hotspots of high host diversity will also result conservation of high specialist parasite diversity.

Changes in community composition across spaces is characterized by β diversity. Modeling β diversity is thus an important tool for the planning of effective habitat conservation because small extensions of protected areas in regions with high β diversity could significantly increase the number of species conserved, as well as the viability of specialized local populations (Jankowski et al., 2009; McNew et al., 2021). The total β diversity has been partitioned into two additive components attributable to differences in taxon richness (e.g., due to nestedness of lineages) or taxon replacement (e.g., due to lineage turnover) between communities (Cardoso et al., 2013). Baselga et al. (2015) identified that the richness and replacement components of β diversity tend to reflect community responses to deterministic (e.g., non-random extinction due to differences in climate) vs. stochastic (e.g., ecological drift due to geographic isolation). In our study, across all sites, phylogenetic β diversity due to changes in richness was found to be significantly higher in the specialist *Haemoproteus* vs. the generalist *Plasmodium*. This can be expected due to the highly nested community structures of specialist species (Dos Anjos et al., 2019) where in each site, the most specialized species would be lost to disturbance (McKinney & Lockwood, 1999; Olden et al., 2004; Devictor et al., 2007; Clavel et al., 2011). We also find that across all sites, phylogenetic β diversity due to taxon replacement was significantly lower in *Haemoproteus* vs. *Plasmodium*. This is because the narrow niche-breadths of specialist species do not allow one specialist to easily replace another, while the broader niches of generalists allow easy replacement of another between sites (Clavel et al., 2011; Carscadden et al., 2020).

Indeed, the relative importance of richness (nestedness) vs. replacement (turnover) has been proposed to be an effective index of generalism, with specialists showing high nestedness and generalists high turnover (Morley et al., 2024).

With regards to the specific variables that predicted *Haemoproteus* phylogenetic β diversity, host functional α diversity and maximum temperature of the warmest month (BIO05) were the most important predictors, followed by the variables related to proportion of grasslands and plantations. In contrast, the most important predictors of *Plasmodium* phylogenetic β diversity were the proportion of degraded habitat, followed closely by isothermality and geographic distance. Importantly, unlike in *Haemoproteus*, host ecology was not found to be a predictor of *Plasmodium* β diversity, emphasizing that generalist species are less affected by host factors than specialists. Critically, the predicted spatial patterns of *Haemoproteus* β diversity indicate that even parasite communities separated by large geographic distances, such as communities separated by the Palghat gap, an ancient (500 Mya) and wide (maximum 40 km) biogeographic gap (Robin et al., 2010), can be very similar to each other, if their bird communities are similar. On the other hand, predicted spatial patterns of *Plasmodium* β diversity indicate that communities separated by larger geographic distances are more likely to be dissimilar from each other. Two distinct clusters—one primarily north of the Palghat gap, and the other primarily south of it—highlight this pattern. Classical literature has considered how hosts may be analogous to islands from the perspective of parasite colonization (Kuris et al., 1980). Our data indicates that in the specialist parasite (*Haemoproteus*), host species can be considered islands as host phylogenetic and ecological differences shape parasite community structure. However, in specialist parasite (*Plasmodium*) spatially distinct host communities serve as islands. This difference between the parasite taxa explains why a strong concordance is found between drivers of parasite β diversity and host functional and phylogenetic β diversity in the specialist *Haemoproteus* but not the generalist *Plasmodium*.

To conclude, we used avian haemosporidian parasites as a model system to identify factors affecting parasite diversity patterns and test if these patterns differ between specialist (*Haemoproteus*) and generalist (*Plasmodium*) taxa. As hypothesized, the drivers of diversity differ in generalist versus specialist parasites. By examining avian haemosporidian parasites, our findings reveal that specialist parasites exhibit higher phylogenetic α diversity, primarily influenced by host-specific factors, while generalist parasites show greater α evenness and are shaped by a broader array of variables, including both host and environmental factors. Specialists are also found to have greater β diversity related to richness likely because specialists tend to be lost without concomitant replacement, leading to a nested community structure in these taxa. In contrast, generalists display greater β diversity related to replacement, as their broad niche-breadth allows them to easily replace one another across the landscape. Similar to α diversity, specialist β diversity was primarily driven by host related factors, while generalist β diversity was driven by a wider variety of non-host related factors. Importantly, as the drivers of diversity patterns differ between specialist and generalist parasites, so do the consequences of their loss.

Critically, specialist parasites seem to be good indicators of ecosystem health because, compared to generalists, specialist parasites are more closely linked to specific hosts, more prone to co-extinctions, and hence more sensitive to disturbance (Rezende et al., 2007; Dunn et al., 2009; Thompson et al., 2018; Carlson et al., 2020). Generalist parasites have broader niche-breadths than specialist parasites, and are thus more likely to be associated with rapid shifts to novel and naive host species, and hence with EIDs such as the chytrid fungus in amphibians (Pounds et al., 2006; Skerratt et al., 2007) and the white nose syndrome in bats (Blehert et al., 2009).

## Data Accessibility Statement

The raw data relating to bird and parasite samples were obtained from Gupta et al. (2019), and are available at https://doi.org/10.6084/m9.figshare.c.4509134.v2. The, bird and parasite genetic data have been deposited at the National Center for Biotechnology Information and GenBank Accession Numbers are reported in Robin et al. (2015) and Gupta et al. (2019), respectively. The bird functional traits were obtained from the Avonet database (https://opentraits.org/datasets/avonet). Environmental data were sourced from the following locations: canopy height (http://lidarradar.jpl.nasa.gov), climate (http://chelsa-climate.org/bioclim), elevation (http://www.earthenv.org/DEM), water flow accumulation (https://www.hydrosheds.org). Landcover data was sourced directly from the authors of Roy et al. (2015). Additionally, we also obtained data on protected area boundaries (http://datasets.wri.org) and the Indian human population (http://www.ciesin.columbia.edu/data/india-census-grids). The code and data used for the analyses reported in this study are available at: URL HERE.

## Supporting Information

Additional supporting information can be found online in the Supporting Information section

## Acknowledgements

We thank the Forest Departments, particularly then PCCFs, of Kerala (R. Rajaraja Varma, V. Gopinathan, T. M. Manoharan and B. S. Corrie) and Tamil Nadu (R. Sunderaraju) for bird sampling permits. We also thank the CCFs, DFOs and range officers who helped with field logistics. We thank our long-term collaborators C. K. Vishnudas, Pooja Gupta, Uma Ramakrishnan and V. V. Robin for all their help and support. We acknowledge financial support from Krea University through a Krea Research Grant to G.D.

## Author Contributions

G.D. conceived the study and performed data analyses; D.J., T.K. and N.N wrote the first draft of the paper. All authors revised and edited the manuscript.

## References

Alexander, N., & Wint, W. (2013). Projected Population Proximity Indices (30km) for 2005, 2030 and 2050. Journal of Open Public Health Data, 1(1). doi:10.5334/jophd.ab

Arellano, L., & Halffter, G. (2022). Gamma diversity: derived from and a determinant of alpha diversity and beta diversity. An analysis of three tropical landscapes. Acta Zool. Mex.(90), 27–76. doi:10.21829/azm.2003.902550

Asghar, M., Hasselquist, D., Hansson, B., Zehtindjiev, P., Westerdahl, H., & Bensch, S. (2015). Hidden costs of infection: Chronic malaria accelerates telomere degradation and senescence in wild birds. Science, 347(6220), 436–438. doi:10.1126/science.1261121

Atkinson, C. T., & Samuel, M. D. (2010). Avian malaria Plasmodium relictum in native Hawaiian forest birds: epizootiology and demographic impacts on LJapapane Himatione sanguinea. Journal of Avian Biology, 41(4), 357–366. doi:10.1111/j.1600-048X.2009.04915.x

Baselga, A., Leprieur, F., & Spencer, M. (2015). Comparing methods to separate components of beta diversity. Methods in Ecology and Evolution, 6(9), 1069–1079. doi:10.1111/2041-210x.12388

Beadell, J. S., Covas, R., Gebhard, C., Ishtiaq, F., Melo, M., Schmidt, B. K., … Fleischer, R. C. (2009). Host associations and evolutionary relationships of avian blood parasites from West Africa. Int J Parasitol, 39(2), 257–266. doi:10.1016/j.ijpara.2008.06.005

Beadell, J. S., Gering, E., Austin, J., Dumbacher, J. P., Peirce, M. A., Pratt, T. K., … Fleischer, R. C. (2004). Prevalence and differential host-specificity of two avian blood parasite genera in the Australo-Papuan region. Mol Ecol, 13(12), 3829–3844. doi:10.1111/j.1365-294X.2004.02363.x

Bensch, S., Hellgren, O., & Pérez-Tris, J. (2009). MalAvi: a public database of malaria parasites and related haemosporidians in avian hosts based on mitochondrial cytochrome b lineages. Mol. Ecol. Resour., 9(5), 1353–1358. doi:10.1111/j.1755-0998.2009.02692.x

Bensch, S., Pérez-Tris, J., Waldenström, J., & Hellgren, O. (2004). Linkage between nuclear and mitochondrial DNA sequences in avian malaria parasites: multiple cases of cryptic speciation? Evolution, 58(7), 1617–1621. doi:10.1111/j.0014-3820.2004.tb01742.x

Betts, A., Gray, C., Zelek, M., MacLean, R. C., & King, K. C. (2018). High parasite diversity accelerates host adaptation and diversification. Science, 360(6391), 907–911. doi:10.1126/science.aam9974

Blehert, D. S., Hicks, A. C., Behr, M., Meteyer, C. U., Berlowski-Zier, B. M., Buckles, E. L., … Stone, W. B. (2009). Bat white-nose syndrome: An emerging fungal pathogen? Science, 323(5911), 227. doi:10.1126/science.1163874

Boyles, J. G., & Storm, J. J. (2007). The perils of picky eating: dietary breadth is related to extinction risk in insectivorous bats. PLoS One, 2(7), e672. doi:10.1371/journal.pone.0000672

Brian, J. I. (2023). Parasites in biodiversity conservation: friend or foe? Trends Parasitol, 39(8), 618–621. doi:10.1016/j.pt.2023.05.005

Brian, J. I., & Aldridge, D. C. (2023). Factors at multiple scales drive parasite community structure. J. Anim. Ecol., 92(2), 377–390. doi:10.1111/1365-2656.13853

Brian, J. I., & Aldridge, D. C. (2024). Host and parasite identity interact in scale-dependent fashion to determine parasite community structure. Oecologia, 204(1), 199–211. doi:10.1007/s00442-023-05499-3

Calcagno, V., Jarne, P., Loreau, M., Mouquet, N., & David, P. (2017). Diversity spurs diversification in ecological communities. Nat. Commun., 8(1), 15810. doi:10.1038/ncomms15810

Cardillo, M., Dinnage, R., & McAlister, W. (2018). The relationship between environmental niche breadth and geographic range size across plant species. Journal of Biogeography, 46(1), 97–109. doi:10.1111/jbi.13477

Cardoso, P., Rigal, F., Carvalho, J. C., Fortelius, M., Borges, P. A. V., Podani, J., … Veech, J. (2013). Partitioning taxon, phylogenetic and functional beta diversity into replacement and richness difference components. Journal of Biogeography, 41(4), 749–761. doi:10.1111/jbi.12239

Cardoso, P., Rigal, F., Carvalho, J. C., & Kembel, S. (2014). BAT -Biodiversity Assessment Tools, an R package for the measurement and estimation of alpha and beta taxon, phylogenetic and functional diversity. Methods in Ecology and Evolution, 6(2), 232–236. doi:10.1111/2041-210x.12310

Carlson, C. J., Dallas, T. A., Alexander, L. W., Phelan, A. L., & Phillips, A. J. (2020). What would it take to describe the global diversity of parasites? Proc. Biol. Sci., 287(1939), 20201841. doi:10.1098/rspb.2020.1841

Carrara, E., Arroyo-Rodríguez, V., Vega-Rivera, J. H., Schondube, J. E., de Freitas, S. M., & Fahrig, L. (2015). Impact of landscape composition and configuration on forest specialist and generalist bird species in the fragmented Lacandona rainforest, Mexico. Biological Conservation, 184, 117–126. doi:10.1016/j.biocon.2015.01.014

Carscadden, K. A., Emery, N. C., Arnillas, C. A., Cadotte, M. W., Afkhami, M. E., Gravel, D., … Wiens, J. J. (2020). Niche Breadth: Causes and Consequences for Ecology, Evolution, and Conservation. The Quarterly Review of Biology, 95(3), 179–214. doi:10.1086/710388

Ceballos, G., Ehrlich, P. R., Barnosky, A. D., García, A., Pringle, R. M., & Palmer, T. M. (2015). Accelerated modern human-induced species losses: Entering the sixth mass extinction. Sci. Adv., 1(5), e1400253. doi:10.1126/sciadv.1400253

Cebrian-Camison, S., Martinez-de la Puente, J., Ruiz-Lopez, M. J., & Figuerola, J. (2025). Do specialist and generalist parasites differ in their prevalence and intensity of infection? A test of the niche breadth and trade-off hypotheses. Int J Parasitol, 55(2), 129–136. doi:10.1016/j.ijpara.2024.11.009

Clavel, J., Julliard, R., & Devictor, V. (2011). Worldwide decline of specialist species: toward a global functional homogenization? Front. Ecol. Environ., 9(4), 222–228. doi:10.1890/080216

Cloyed, C. S., Balmer, B. C., Schwacke, L. H., Wells, R. S., Berens McCabe, E. J., Barleycorn, A. A., … Carmichael, R. H. (2021). Interaction between dietary and habitat niche breadth influences cetacean vulnerability to environmental disturbance. Ecosphere, 12(9). doi:10.1002/ecs2.3759

Cooper, N., Griffin, R., Franz, M., Omotayo, M., Nunn, C. L., & Fryxell, J. (2012). Phylogenetic host specificity and understanding parasite sharing in primates. Ecol Lett, 15(12), 1370–1377. doi:10.1111/j.1461-0248.2012.01858.x

Devictor, V., Julliard, R., Clavel, J., Jiguet, F., Lee, A., & Couvet, D. (2008). Functional biotic homogenization of bird communities in disturbed landscapes. Global Ecology and Biogeography: A Journal of Macroecology, 17(2), 252–261. doi:10.1111/j.1466-8238.2007.00364.x

Devictor, V., Julliard, R., Couvet, D., Lee, A., & Jiguet, F. (2007). Functional homogenization effect of urbanization on bird communities. Conserv. Biol., 21(3), 741–751. doi:10.1111/j.1523-1739.2007.00671.x

Dharmarajan, G., Gupta, P., Vishnudas, C. K., & Robin, V. V. (2021). Anthropogenic disturbance favours generalist over specialist parasites in bird communities: Implications for risk of disease emergence. Ecology Letters, 24(9), 1859–1868. doi:10.1111/ele.13818

Dharmarajan, G., Li, R., Chanda, E., Dean, K. R., Dirzo, R., Jakobsen, K. S., … Stenseth, N. C. (2022). The animal origin of major human infectious diseases: What can past epidemics teach us about preventing the next pandemic? Zoonoses, 2, 11. doi:10.15212/zoonoses-2021-0028

Dimitrov, D., Zehtindjiev, P., & Bensch, S. (2010). Genetic diversity of avian blood parasites in SE Europe: Cytochrome b lineages of the genera Plasmodium and Haemoproteus (Haemosporida) from Bulgaria. Acta Parasitologica, 55(3). doi:10.2478/s11686-010-0029-z

Dixon, P. (2003). VEGAN, a package of R functions for community ecology. Journal of Vegetation Science, 14(6), 927–930. doi:10.1111/j.1654-1103.2003.tb02228.x

Dobson, A., Lafferty, K. D., Kuris, A. M., Hechinger, R. F., & Jetz, W. (2008). Homage to Linnaeus: How many parasites? How many hosts? Proceedings of the National Academy of Sciences of the United States of America, 105 Suppl 1, 11482–11489. doi:10.1073/pnas.0803232105

Dos Anjos, L., Bochio, G. M., Medeiros, H. R., Almeida, B. d. A., Lindsey, B. R. A., Calsavara, L. C., … Domingues Torezan, J. M. (2019). Insights on the functional composition of specialist and generalist birds throughout continuous and fragmented forests. Ecology and Evolution, 9(11), 6318–6328. doi:10.1002/ece3.5204

Doussang, D., Sallaberry-Pincheira, N., Cabanne, G. S., Lijtmaer, D. A., Gonzalez-Acuna, D., & Vianna, J. A. (2021). Specialist versus generalist parasites: the interactions between host diversity, environment and geographic barriers in avian malaria. Int J Parasitol, 51(11), 899–911. doi:10.1016/j.ijpara.2021.04.003

Dunn, R. R., Harris, N. C., Colwell, R. K., Koh, L. P., & Sodhi, N. S. (2009). The sixth mass coextinction: Are most endangered species parasites and mutualists? Proceedings of the Royal Society of London. Series B: Biological Sciences, 276(1670), 3037–3045. doi:10.1098/rspb.2009.0413

Dutra, D. d. A. (2023). Revealing the drivers of parasite diversity: territorial and biodiverse hosts raise haemosporidian regional diversity worldwide. doi:10.22541/au.168232922.26436194/v1

Ebert, D. (2025). The Red Queen and the Timescale of Antagonistic Coevolution: Parasite Selection for Genetic Diversity. Annu Rev Genet, 59(1), 215–236. doi:10.1146/annurev-genet-112024-091229

Fallon, S. M., Bermingham, E., & Ricklefs, R. E. (2005). Host specialization and geographic localization of avian malaria parasites: a regional analysis in the Lesser Antilles. Am Nat, 165(4), 466–480. doi:10.1086/428430

Farrell, M. J., Berrang-Ford, L., & Davies, T. J. (2013). The study of parasite sharing for surveillance of zoonotic diseases. Environmental Research Letters, 8(1). doi:Artn 015036 10.1088/1748-9326/8/1/015036

Fenton, A., & Brockhurst, M. A. (2008). The role of specialist parasites in structuring host communities. Ecol. Res., 23(5), 795–804. doi:10.1007/s11284-007-0440-6

Ferrier, S., Manion, G., Elith, J., & Richardson, K. (2007). Using generalized dissimilarity modelling to analyse and predict patterns of beta diversity in regional biodiversity assessment. Diversity and Distributions, 13(3), 252–264. doi:10.1111/j.1472-4642.2007.00341.x

Fitzpatrick, M. C., Mokany, K., Manion, G., Nieto-Lugilde, D., & Ferrier, S. (2021). gdm: Generalized dissimilarity modeling. R package version1.5.0-1. https://CRAN.R-project.org/package=gdm.

Frainer, A., McKie, B. G., Amundsen, P. A., Knudsen, R., & Lafferty, K. D. (2018). Parasitism and the biodiversity-functioning relationship. Trends in Ecology and Evolution, 33(4), 260–268. doi:10.1016/j.tree.2018.01.011

Freeman, M. T., Olivier, P. I., & van Aarde, R. J. (2018). Matrix transformation alters species-area relationships in fragmented coastal forests. Landscape Ecology, 33(2), 307–322. doi:10.1007/s10980-017-0604-x

Gibb, R., Redding, D. W., Chin, K. Q., Donnelly, C. A., Blackburn, T. M., Newbold, T., & Jones, K. E. (2020). Zoonotic host diversity increases in human-dominated ecosystems. Nature. doi:10.1038/s41586-020-2562-8

Goyal, N., Sharma, A., Jagati, V., Herur, A., Jain, A., Gopal, A., … Robin, V. V. (2025). Spatial Patterns of Diversity in Forest Birds of Peninsular India. Ecology and Evolution, 15(12), e72698. doi:10.1002/ece3.72698

Granot, I., & Belmaker, J. (2020). Niche breadth and species richness: Correlation strength, scale and mechanisms. Global Ecology and Biogeography: A Journal of Macroecology, 29(1), 159–170. doi:10.1111/geb.13011

Gupta, P., Robin, V. V., & Dharmarajan, G. (2020). Towards a more healthy conservation paradigm: integrating disease and molecular ecology to aid biological conservation. Journal of Genetics, 99(1), Article 65, 61–26. doi:10.1007/s12041-020-01225-7

Gupta, P., Vishnudas, C. K., Ramakrishnan, U., Robin, V. V., & Dharmarajan, G. (2019). Geographical and host species barriers differentially affect generalist and specialist parasite community structure in a tropical sky-island archipelago. Proceedings of the Royal Society B, 286(1904), 20190439. doi:10.1098/rspb.2019.0439

Hartley, S., & Kunin, W. E. (2003). Scale dependency of rarity, extinction risk, and conservation priority. Conserv. Biol., 17(6), 1559–1570. doi:10.1111/j.1523-1739.2003.00015.x

Hatcher, M. J., Dick, J. T. A., & Dunn, A. M. (2012a). Disease emergence and invasions. Functional Ecology, 26(6), 1275–1287. doi:10.1111/j.1365-2435.2012.02031.x

Hatcher, M. J., Dick, J. T. A., & Dunn, A. M. (2012b). Diverse effects of parasites in ecosystems: linking interdependent processes. Front. Ecol. Environ., 10(4), 186–194. doi:10.1890/110016

Hellgren, O., Bensch, S., & Malmqvist, B. (2008). Bird hosts, blood parasites and their vectors--associations uncovered by molecular analyses of blackfly blood meals. Molecular Ecology, 17(6), 1605–1613. doi:10.1111/j.1365-294X.2007.03680.x

Hellgren, O., Waldenstrom, J., & Bensch, S. (2004). A new PCR assay for simultaneous studies of Leucocytozoon, Plasmodium, and Haemoproteus from avian blood. J Parasitol, 90(4), 797–802. doi:10.1645/GE-184R1

Hudson, P. J., Dobson, A. P., & Lafferty, K. D. (2006). Is a healthy ecosystem one that is rich in parasites? Trends in Ecology and Evolution, 21(7), 381–385. doi:10.1016/j.tree.2006.04.007

Ibarra, J. T., & Martin, K. (2015). Biotic homogenization: Loss of avian functional richness and habitat specialists in disturbed Andean temperate forests. Biological Conservation, 192, 418–427. doi:10.1016/j.biocon.2015.11.008

IlgūBnas, M., Bukauskaitė, D., Palinauskas, V., Iezhova, T. A., Dinhopl, N., Nedorost, N., … ValkiūBnas, G. (2016). Mortality and pathology in birds due to Plasmodium (Giovannolaia) homocircumflexum infection, with emphasis on the exoerythrocytic development of avian malaria parasites. Malar. J., 15(1), 256. doi:10.1186/s12936-016-1310-x

Jacob, S. T., Crozier, I., Fischer, W. A., 2nd, Hewlett, A., Kraft, C. S., Vega, M. A., … Kuhn, J. H. (2020). Ebola virus disease. Nat Rev Dis Primers, 6(1), 13. doi:10.1038/s41572-020-0147-3

Jankowski, J. E., Ciecka, A. L., Meyer, N. Y., & Rabenold, K. N. (2009). Beta diversity along environmental gradients: implications of habitat specialization in tropical montane landscapes. J. Anim. Ecol., 78(2), 315–327. doi:10.1111/j.1365-2656.2008.01487.x

Johnson, P. T., de Roode, J. C., & Fenton, A. (2015). Why infectious disease research needs community ecology. Science, 349(6252), 1259504. doi:10.1126/science.1259504

Kamiya, T., O’Dwyer, K., Nakagawa, S., & Poulin, R. (2014). Host diversity drives parasite diversity: metaLJanalytical insights into patterns and causal mechanisms. Ecography (Cop.), 37(7), 689–697. doi:10.1111/j.1600-0587.2013.00571.x

Kearse, M., Moir, R., Wilson, A., Stones-Havas, S., Cheung, M., Sturrock, S., … Duran, C. (2012). Geneious Basic: an integrated and extendable desktop software platform for the organization and analysis of sequence data. Bioinformatics, 28(12), 1647–1649.

Keesing, F., & Ostfeld, R. S. (2021). Impacts of biodiversity and biodiversity loss on zoonotic diseases. Proc. Natl. Acad. Sci. U. S. A., 118(17), e2023540118. doi:10.1073/pnas.2023540118

Kotiaho, J. S., Kaitala, V., Komonen, A., & Päivinen, J. (2005). Predicting the risk of extinction from shared ecological characteristics. Proc Natl Acad Sci U S A, 102(6), 1963–1967. doi:10.1073/pnas.0406718102

Kuhn, M. (2008). Building predictive models in R using the caret package. Journal of Statistical Software, 28(5), 1–26. doi:10.18637/jss.v028.i05

Kuris, A. M., Blaustein, A. R., & Alio, J. J. (1980). Hosts as islands. The American Naturalist, 116(4), 570–586.

Lafferty, K. D. (2012). Biodiversity loss decreases parasite diversity: theory and patterns. Philos. Trans. R. Soc. Lond. B Biol. Sci., 367(1604), 2814–2827. doi:10.1098/rstb.2012.0110

Lesica, P., Yurkewycz, R., & Crone, E. E. (2006). Rare plants are common where you find them. Am. J. Bot., 93(3), 454–459. doi:10.3732/ajb.93.3.454

Liang, C., Yang, G., Wang, N., Feng, G., Yang, F., Svenning, J.-C., & Yang, J. (2019). Taxonomic, phylogenetic and functional homogenization of bird communities due to land use change. Biological Conservation, 236, 37–43. doi:10.1016/j.biocon.2019.05.036

Lôbo, D., Leão, T., Melo, F. P. L., Santos, A. M. M., & Tabarelli, M. (2011). Forest fragmentation drives Atlantic forest of northeastern Brazil to biotic homogenization. Diversity and Distributions, 17(2), 287–296. doi:10.1111/j.1472-4642.2010.00739.x

Lorel, C., Le Viol, I., Plutzar, C., Jiguet, F., & Mouchet, M. (2021). Linking the diversity and structure of French avian communities with landscape parameters, climate and NPP flows. Regional Environmental Change, 21(2). doi:10.1007/s10113-021-01786-y

Lymbery, A. J., & Smit, N. J. (2023). Conservation of parasites: A primer. Int J Parasitol Parasites Wildl, 21, 255–263. doi:10.1016/j.ijppaw.2023.07.001

Macarthur, R., & Levins, R. (1967). The limiting similarity, convergence, and divergence of coexisting species. Am. Nat., 101(921), 377–385. doi:10.1086/282505

MacArthur, R. H. (1965). Patterns of species diversity. Biological reviews, 40(4), 510–533.

Maechler, M., Rousseeuw, P., Struyf, A., Hubert, M., & Hornik, K. (2023). cluster: Cluster Analysis Basics and Extensions. R package version 2.1.6.

Manzoli, D. E., Saravia-Pietropaolo, M. J., Arce, S. I., Percara, A., Antoniazzi, L. R., & Beldomenico, P. M. (2021). Specialist by preference, generalist by need: availability of quality hosts drives parasite choice in a natural multihost-parasite system. Int. J. Parasitol., 51(7), 527–534. doi:10.1016/j.ijpara.2020.12.003

Marcogliese, D. J. (2005). Parasites of the superorganism: Are they indicators of ecosystem health? International Journal for Parasitology, 35(7), 705–716. doi:10.1016/j.ijpara.2005.01.015

Martins, P. M., Poulin, R., & Goncalves-Souza, T. (2021). Drivers of parasite beta-diversity among anuran hosts depend on scale, realm and parasite group. Philos Trans R Soc Lond B Biol Sci, 376(1837), 20200367. doi:10.1098/rstb.2020.0367

McKinney, M. L., & Lockwood, J. L. (1999). Biotic homogenization: a few winners replacing many losers in the next mass extinction. Trends in Ecology & Evolution, 14(11), 450–453. doi:10.1016/s0169-5347(99)01679-1

McNew, S. M., Barrow, L. N., Williamson, J. L., Galen, S. C., Skeen, H. R., DuBay, S. G., … Witt, C. C. (2021). Contrasting drivers of diversity in hosts and parasites across the tropical Andes. Proc. Natl. Acad. Sci. U. S. A., 118(12), e2010714118. doi:10.1073/pnas.2010714118

Moens, M. A., & Perez-Tris, J. (2016). Discovering potential sources of emerging pathogens: South America is a reservoir of generalist avian blood parasites. Int J Parasitol, 46(1), 41–49. doi:10.1016/j.ijpara.2015.08.001

Moens, M. A., Valkiunas, G., Paca, A., Bonaccorso, E., Aguirre, N., & Perez-Tris, J. (2016). Parasite specialization in a unique habitat: hummingbirds as reservoirs of generalist blood parasites of Andean birds. J Anim Ecol, 85(5), 1234–1245. doi:10.1111/1365-2656.12550

Mokany, K., Ware, C., Woolley, S. N. C., Ferrier, S., Fitzpatrick, Matthew C., & Bahn, V. (2022). A working guide to harnessing generalized dissimilarity modelling for biodiversity analysis and conservation assessment. Global Ecology and Biogeography, 31(4), 802–821. doi:10.1111/geb.13459

Moore, J. H., Gibson, L., Amir, Z., Chanthorn, W., Ahmad, A. H., Jansen, P. A., … Luskin, M. S. (2023). The rise of hyperabundant native generalists threatens both humans and nature. Biol. Rev. Camb. Philos. Soc., 98(5), 1829–1844. doi:10.1111/brv.12985

Morand, S., & Poulin, R. (2000). Nematode parasite species richness and the evolution of spleen size in birds. Canadian Journal of Zoology, 78(8), 1356–1360. doi:10.1139/z00-076

Morley, L., Crain, B. J., Krupnick, G., & Spalink, D. (2024). Turnover importance: Operationalizing beta diversity to quantify the generalism continuum. Methods in Ecology and Evolution, 15(5), 951–964. doi:10.1111/2041-210x.14324

Mougi, A. (2022). Infected food web and ecological stability. Sci. Rep., 12(1), 8139. doi:10.1038/s41598-022-11968-1

Moulatlet, G. M., Merow, C., Maitner, B., Boyle, B., Feng, X., Frazier, A. E., … Enquist, B. J. (2025). General laws of biodiversity: Climatic niches predict plant range size and ecological dominance globally. Proc Natl Acad Sci U S A, 122(46), e2517585122. doi:10.1073/pnas.2517585122

Musa, S., Mackenstedt, U., Woog, F., & Dinkel, A. (2019). Avian malaria on Madagascar: prevalence, biodiversity and specialization of haemosporidian parasites. Int. J. Parasitol., 49(3-4), 199–210. doi:10.1016/j.ijpara.2018.11.001

Ndlovu, M., Wardjomto, M. B., Pori, T., & Nangammbi, T. C. (2024). Diversity and Host Specificity of Avian Haemosporidians in an Afrotropical Conservation Region. Animals (Basel), 14(19). doi:10.3390/ani14192906

Okanga, S., Cumming, G. S., Hockey, P. A., Nupen, L., & Peters, J. L. (2014). Host specificity and co-speciation in avian haemosporidia in the Western Cape, South Africa. PLoS One, 9(2), e86382. doi:10.1371/journal.pone.0086382

Olden, J. D., Douglas, M. E., & Douglas, M. R. (2005). The human dimensions of biotic homogenization. Conserv. Biol., 19(6), 2036–2038. doi:10.1111/j.1523-1739.2005.00288.x

Olden, J. D., Leroy Poff, N., Douglas, M. R., Douglas, M. E., & Fausch, K. D. (2004). Ecological and evolutionary consequences of biotic homogenization. Trends Ecol. Evol., 19(1), 18–24. doi:10.1016/j.tree.2003.09.010

Ostfeld, R. S., & Keesing, F. (2020). Species that can make us ill thrive in human habitats. Nature. doi:10.1038/d41586-020-02189-5

PaganiLJNúñez, E., Liang, D., He, C., Zhou, X., Luo, X., Liu, Y., & Goodale, E. (2019). Niches in the Anthropocene: passerine assemblages show niche expansion from natural to urban habitats. Ecography, 42(8), 1360–1369. doi:10.1111/ecog.04203

Pei, S., Yu, P., Raghwani, J., Wang, Y., Liu, Z., Li, Y., … Tian, H. (2025). Anthropogenic land consolidation intensifies zoonotic host diversity loss and disease transmission in human habitats. Nat Ecol Evol, 9(1), 99–110. doi:10.1038/s41559-024-02570-x

Pellissier, V., Barnagaud, J. Y., Kissling, W. D., Şekercioğlu, Ç., & Svenning, J. C. (2018). Niche packing and expansion account for species richness–productivity relationships in global bird assemblages. Global Ecology and Biogeography, 27(5), 604–615. doi:10.1111/geb.12723

Pigeault, R., Vézilier, J., Cornet, S., Zélé, F., Nicot, A., Perret, P., … Rivero, A. (2015). Avian malaria: a new lease of life for an old experimental model to study the evolutionary ecology of Plasmodium. Philos. Trans. R. Soc. Lond. B Biol. Sci., 370(1675), 20140300. doi:10.1098/rstb.2014.0300

Pigot, Alex L., Trisos, C. H., & Tobias, J. A. (2016). Functional traits reveal the expansion and packing of ecological niche space underlying an elevational diversity gradient in passerine birds. Proc Biol Sci, 283(1822), 20152013. doi:10.1098/rspb.2015.2013

Pinheiro, J. C., & Bates, D. M. (2000). Mixed-Effects Models in S and S-PLUS. New York: Springer.

Pinheiro, J. C., Bates, D. M., & Team, R. C. (2023). nlme: Linear and Nonlinear Mixed Effects Models. R package version 3. 1–164.

Poulin, R., & Morand, S. (2000). The diversity of parasites. Quarterly Review of Biology, 75(3), 277–293. doi:10.1086/393500

Pounds, J. A., Bustamante, M. R., Coloma, L. A., Consuegra, J. A., Fogden, M. P., Foster, P. N., … Young, B. E. (2006). Widespread amphibian extinctions from epidemic disease driven by global warming. Nature, 439(7073), 161–167. doi:10.1038/nature04246

Pyron, R. A., & Pennell, M. (2022). Macroevolutionary perspectives on Anthropocene extinction. Biol. Conserv., 274(109733), Not Available. doi:10.1016/j.biocon.2022.109733

Rezende, E. L., Lavabre, J. E., Guimarães, P. R., Jordano, P., & Bascompte, J. (2007). Non-random coextinctions in phylogenetically structured mutualistic networks. Nature, 448(7156), 925–928. doi:10.1038/nature05956

Robin, V. V., & Nandini, R. (2012). Shola habitats on sky islands: status of research on montane forests and grasslands in southern India. Current Science, 103(12), 1427–1437.

Robin, V. V., Sinha, A., & Ramakrishnan, U. (2010). Ancient geographical gaps and paleo-climate shape the phylogeography of an endemic bird in the sky islands of southern India. PLoS One, 5(10), e13321. doi:10.1371/journal.pone.0013321

Robin, V. V., Vishnudas, C. K., Gupta, P., & Ramakrishnan, U. (2015). Deep and wide valleys drive nested phylogeographic patterns across a montane bird community. Proc Biol Sci, 282(1810). doi:10.1098/rspb.2015.0861

Ronquist, F., & Huelsenbeck, J. P. (2003). MrBayes 3: Bayesian phylogenetic inference under mixed models. Bioinformatics, 19(12), 1572–1574. doi:10.1093/bioinformatics/btg180

Roy, P. S., Behera, M. D., Murthy, M. S. R., Roy, A., Singh, S., Kushwaha, S. P. S., … Ramachandran, R. M. (2015). New vegetation type map of India prepared using satellite remote sensing: Comparison with global vegetation maps and utilities. International Journal of Applied Earth Observation and Geoinformation, 39, 142–159. doi:10.1016/j.jag.2015.03.003

Schoepf, I., Olson, S., Moore, I. T., & Bonier, F. (2022). Experimental reduction of haemosporidian infection affects maternal reproductive investment, parental behaviour and offspring condition. Proc Biol Sci, 289(1987), 20221978. doi:10.1098/rspb.2022.1978

Seutin, G., White, B. N., & Boag, P. T. (1991). Preservation of avian blood and tissue samples for DNA analyses. Canadian Journal of Zoology, 69(1), 82–90. doi:10.1139/z91-013

Skerratt, L. F., Berger, L., Speare, R., Cashins, S., McDonald, K. R., Phillott, A. D., … Kenyon, N. (2007). Spread of Chytridiomycosis has caused the rapid global decline and extinction of frogs. EcoHealth, 4(2), 125–134. doi:10.1007/s10393-007-0093-5

Slatyer, R. A., Hirst, M., & Sexton, J. P. (2013). Niche breadth predicts geographical range size: a general ecological pattern. Ecol Lett, 16(8), 1104–1114. doi:10.1111/ele.12140

Terborgh, J. (2020). At 50, Janzen–Connell has come of age. Bioscience, 70(12), 1082–1092. doi:10.1093/biosci/biaa110

Thieltges, D. W., Hof, C., Dehling, D. M., Brändle, M., Brandl, R., & Poulin, R. (2011). Host diversity and latitude drive trematode diversity patterns in the European freshwater fauna. Global Ecology and Biogeography: A Journal of Macroecology, 20(5), 675–682. doi:10.1111/j.1466-8238.2010.00631.x

Thompson, R. C. A., Lymbery, A. J., & Godfrey, S. S. (2018). Parasites at Risk -Insights from an Endangered Marsupial. Trends Parasitol, 34(1), 12–22. doi:10.1016/j.pt.2017.09.001

Timms, R., & Read, A. F. (1999). What makes a specialist special? Trends in Ecology & Evolution, 14(9), 333–334. doi:10.1016/s0169-5347(99)01697-3

Tobias, J. A., Sheard, C., Pigot, A. L., Devenish, A. J. M., Yang, J., Sayol, F., … Schleuning, M. (2022). AVONET: morphological, ecological and geographical data for all birds. Ecol Lett, 25(3), 581–597. doi:10.1111/ele.13898

Turvey, S. T., & Crees, J. J. (2019). Extinction in the Anthropocene. Curr Biol, 29(19), R982–R986. doi:10.1016/j.cub.2019.07.040

Van Valen, L. (1973). A new evolutionary law. Evolutionary Theory, 1, 1–30.

Verberk, W. C. E. P., van der Velde, G., & Esselink, H. (2010). Explaining abundance-occupancy relationships in specialists and generalists: a case study on aquatic macroinvertebrates in standing waters. J. Anim. Ecol., 79(3), 589–601. doi:10.1111/j.1365-2656.2010.01660.x

Waldenstrom, J., Bensch, S., Kiboi, S., Hasselquist, D., & Ottosson, U. (2002). Cross-species infection of blood parasites between resident and migratory songbirds in Africa. Mol Ecol, 11(8), 1545–1554. doi:10.1046/j.1365-294x.2002.01523.x

Wang, S., Loreau, M., de Mazancourt, C., Isbell, F., Beierkuhnlein, C., Connolly, J., … Craven, D. (2021). Biotic homogenization destabilizes ecosystem functioning by decreasing spatial asynchrony. Ecology, 102(6), e03332. doi:10.1002/ecy.3332

Wells, K., & Clark, N. J. (2019). Host Specificity in Variable Environments. Trends Parasitol, 35(6), 452–465. doi:10.1016/j.pt.2019.04.001

Whittaker, R. H. (1960). Vegetation of the siskiyou mountains, Oregon and California. Ecol. Monogr., 30(3), 279–338. doi:10.2307/1943563

Whittaker, R. H. (1972). Evolution and measurement of species diversity. Taxon, 21(2-3), 213–251. doi:10.2307/1218190

Wright, M. N., & Ziegler, A. (2017). ranger: A fast implementation of random forests for high dimensional data in C++ and R. Journal of Statistical Software, 77(1), 1–17. doi:10.18637/jss.v077.i01

