## Supplementary Material for "Host and environmental factors differentially affect patterns of diversity in specialist and generalist parasites"

**SUPPLEMENTARY MATERIALS**

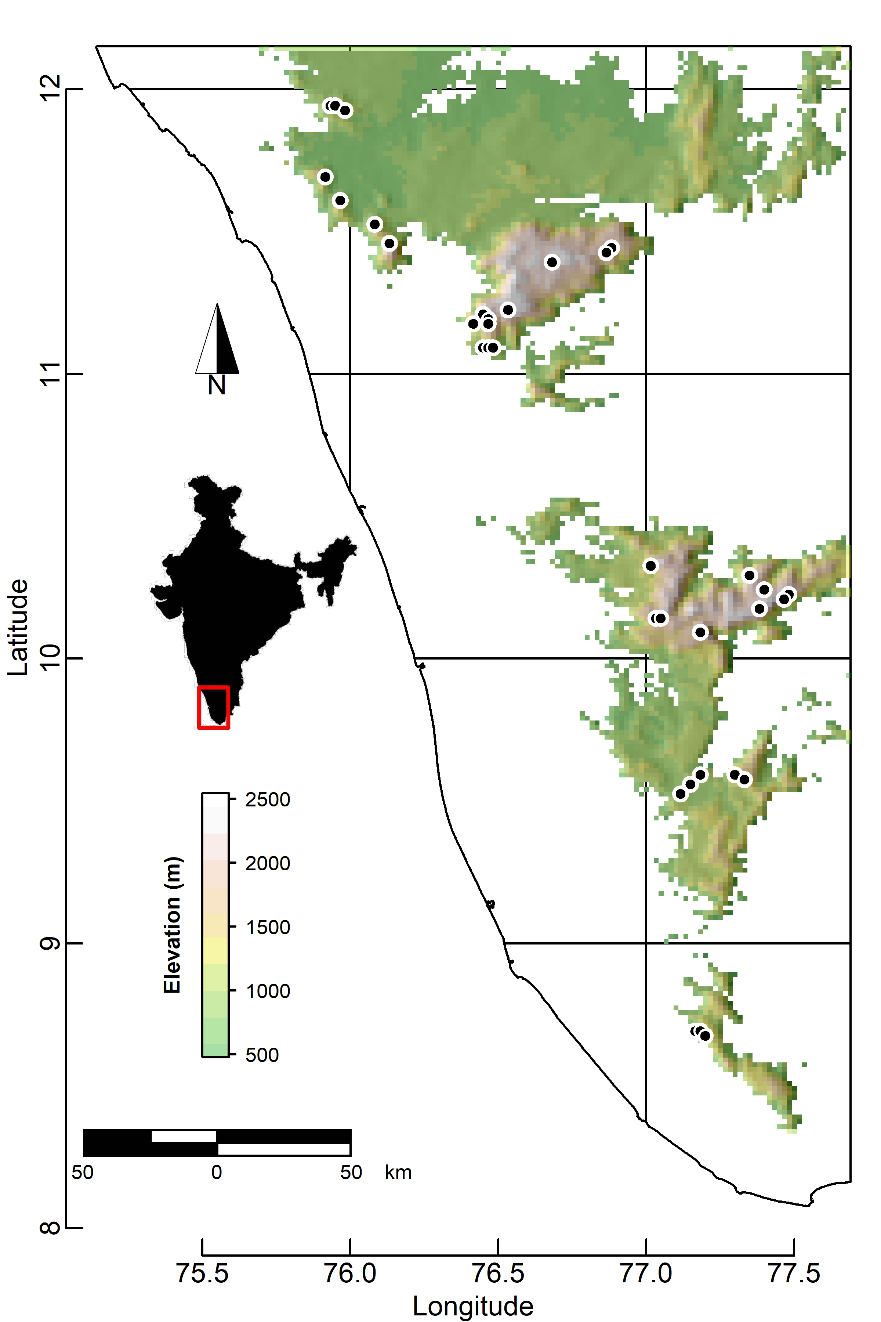

Figure S01: Map of the study area located in the Western Ghats, Southern India, showing the 42 sample site locations (circles) where birds (N = 1172) were sampled (Table S1).

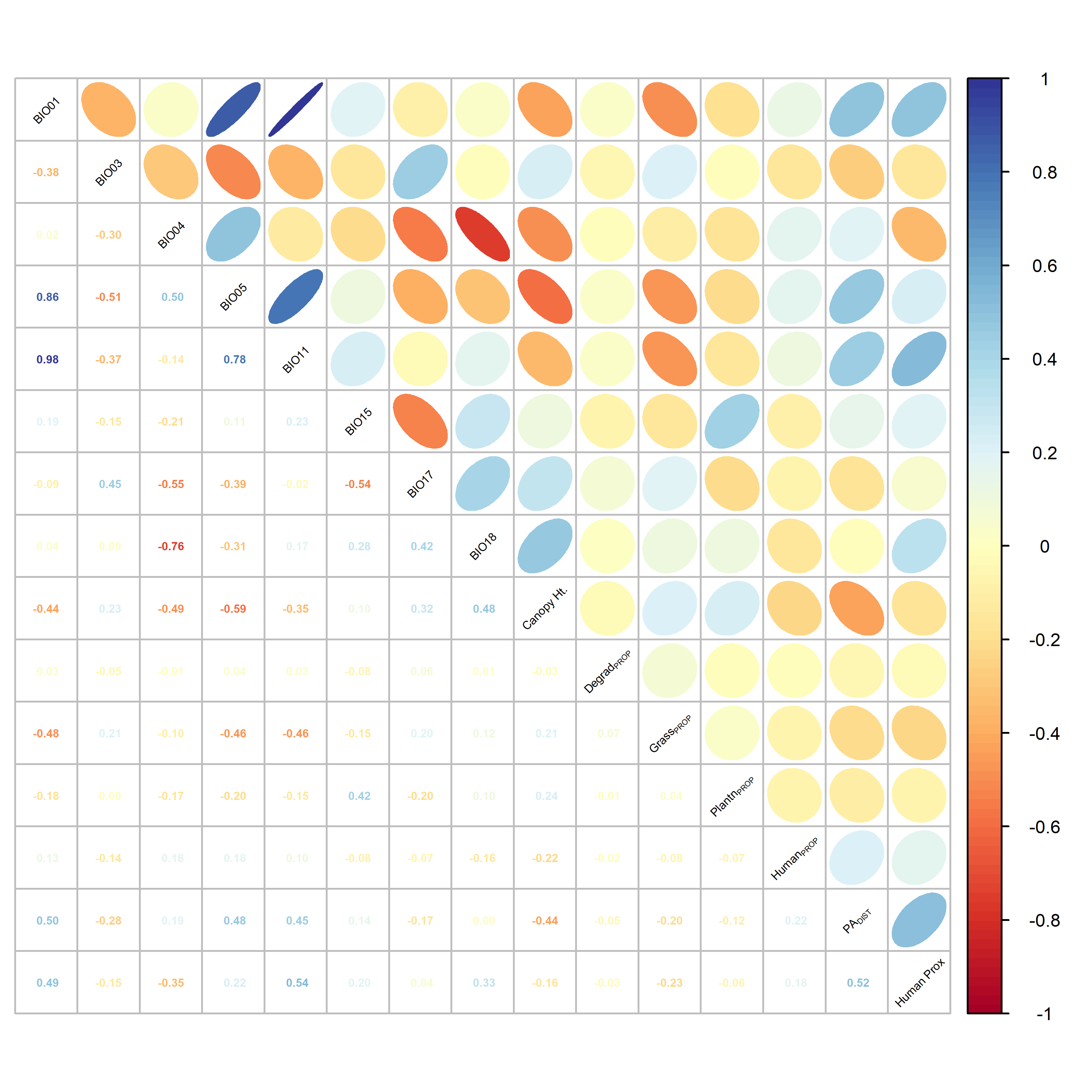

**Figure S02:** Pairwise correlation plot amongst environmental variables used across all final best-fit analytical models. The ellipses (upper triangle) are visual representations of scatter plots, and the numbers (lower triangle) are the Pearson’s correlation coefficients between each pair of variables. Variable names are given in the diagonal, and include: annual mean temperature (BIO01), isothermality (BIO03), temperature seasonality (BIO04), max temperature of warmest month (BIO05), mean temperature of coldest quarter (BIO11), precipitation seasonality (BIO15), precipitation of driest quarter (BIO17), precipitation of warmest quarter (BIO18), canopy height (Canopy-Ht), proportion degraded habitat (Degrad_PROP_), proportion grassland (Grass_PROP_), proportion plantation (Plantn_PROP_), proportion settlements (Human_PROP_), distance to protected areas (PA_DIST_), human proximity index (Human-Prox). Colors range from a perfect negative correlation coefficient (-1; red) to a perfect positive correlation coefficient (1; blue)

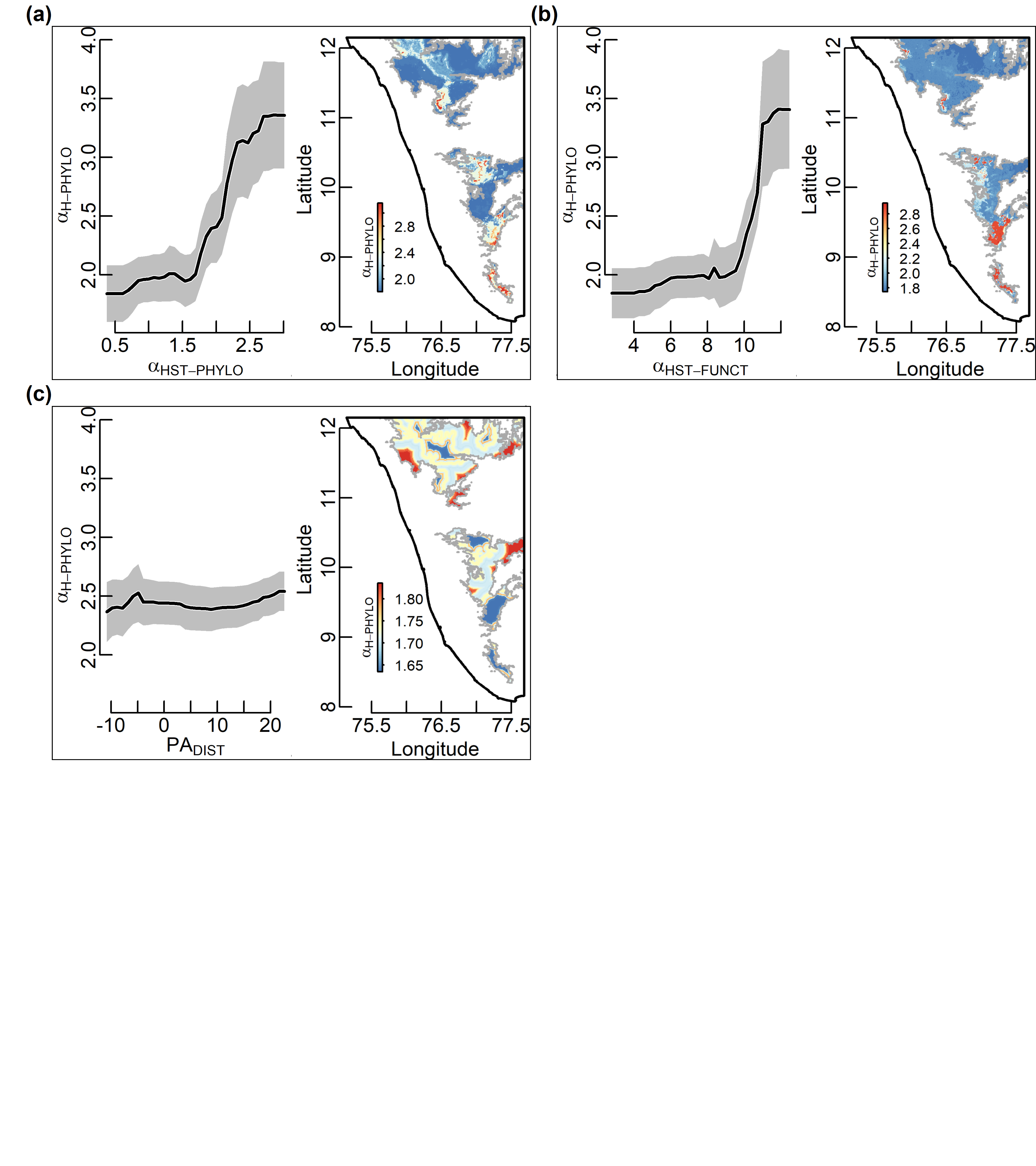

**Figure S03:** Partial dependence plots, based on the final random forest model for *Haemoproteus* phylogenetic α diversity (α_H-PHYLO_) in the Western Ghats, Southern India. Separate figures are plotted for each independent variable in the random forest model, including: (a) host phylogenetic α diversity (α_HST-PHYLO_), (b) host functional α diversity (α_HST-FUNCT_), (c) distance to protected areas (PA_DIST_). In each plot, the figure on the left shows the mean marginal influence of a particular independent variable on α_H-PHYLO_ while holding the other independent variables constant. Alternatively, the figure on the right shows the spatially explicit partial predictions for α_H-PHYLO_ while holding the other independent variables constant.

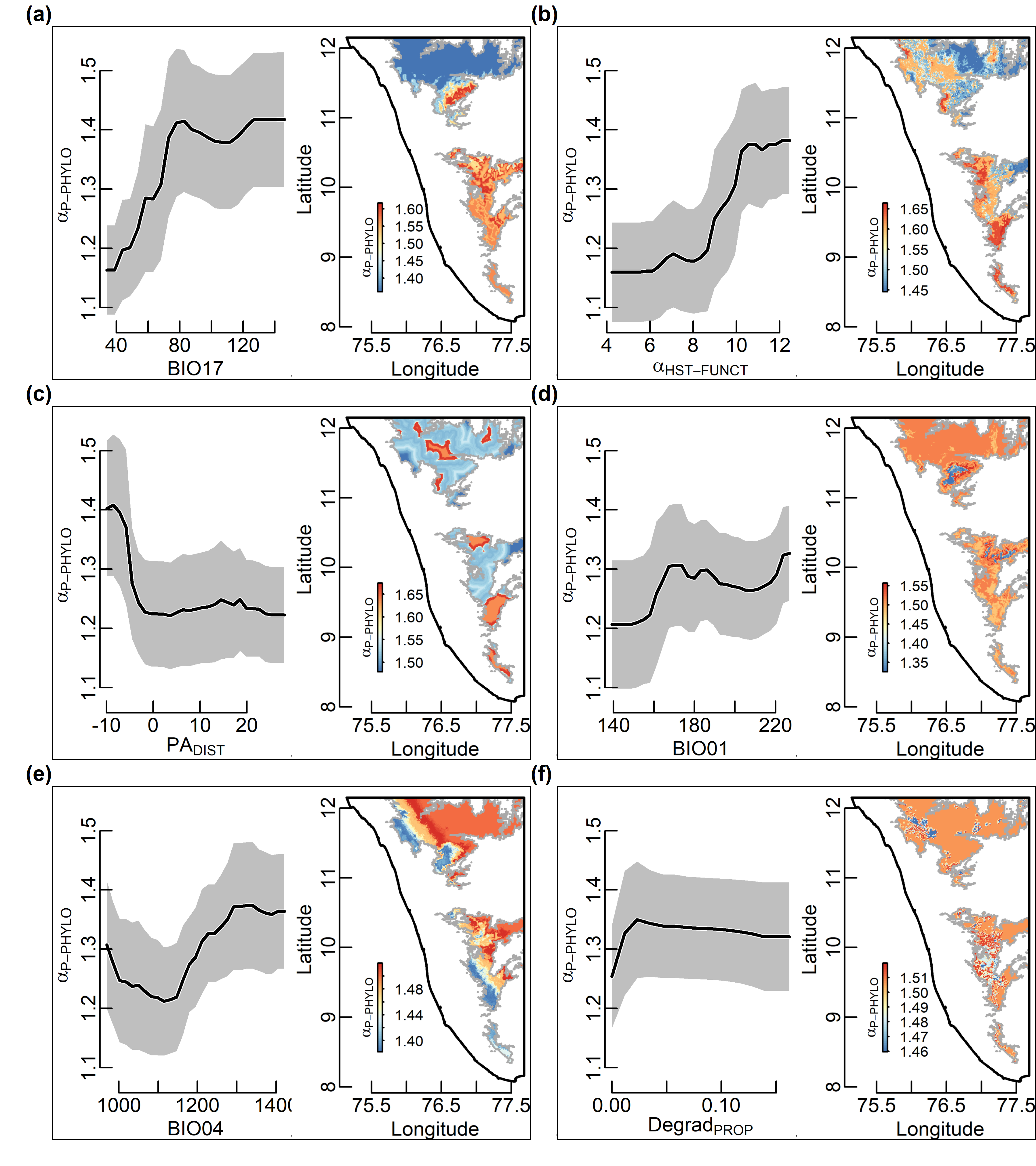

**Figure S04:** Partial dependence plots, based on the final random forest model for *Plasmodium* phylogenetic α diversity (α_P-PHYLO_) in the Western Ghats, Southern India. Separate figures are plotted for each independent variable in the random forest model, including: (a) precipitation of driest quarter (BIO17), (b) host functional α diversity (α_HST-FUNCT_), (c) distance to protected areas (PA_DIST_), (d) annual mean temperature (BIO01), (e) temperature seasonality (BIO04), (f) proportion degraded habitat (Degrad_PROP_). In each plot, the figure on the left shows the mean marginal influence of a particular independent variable on α_P-PHYLO_ while holding the other independent variables constant. Alternatively, the figure on the right shows the spatially explicit partial predictions for α_P-PHYLO_ while holding the other independent variables constant.

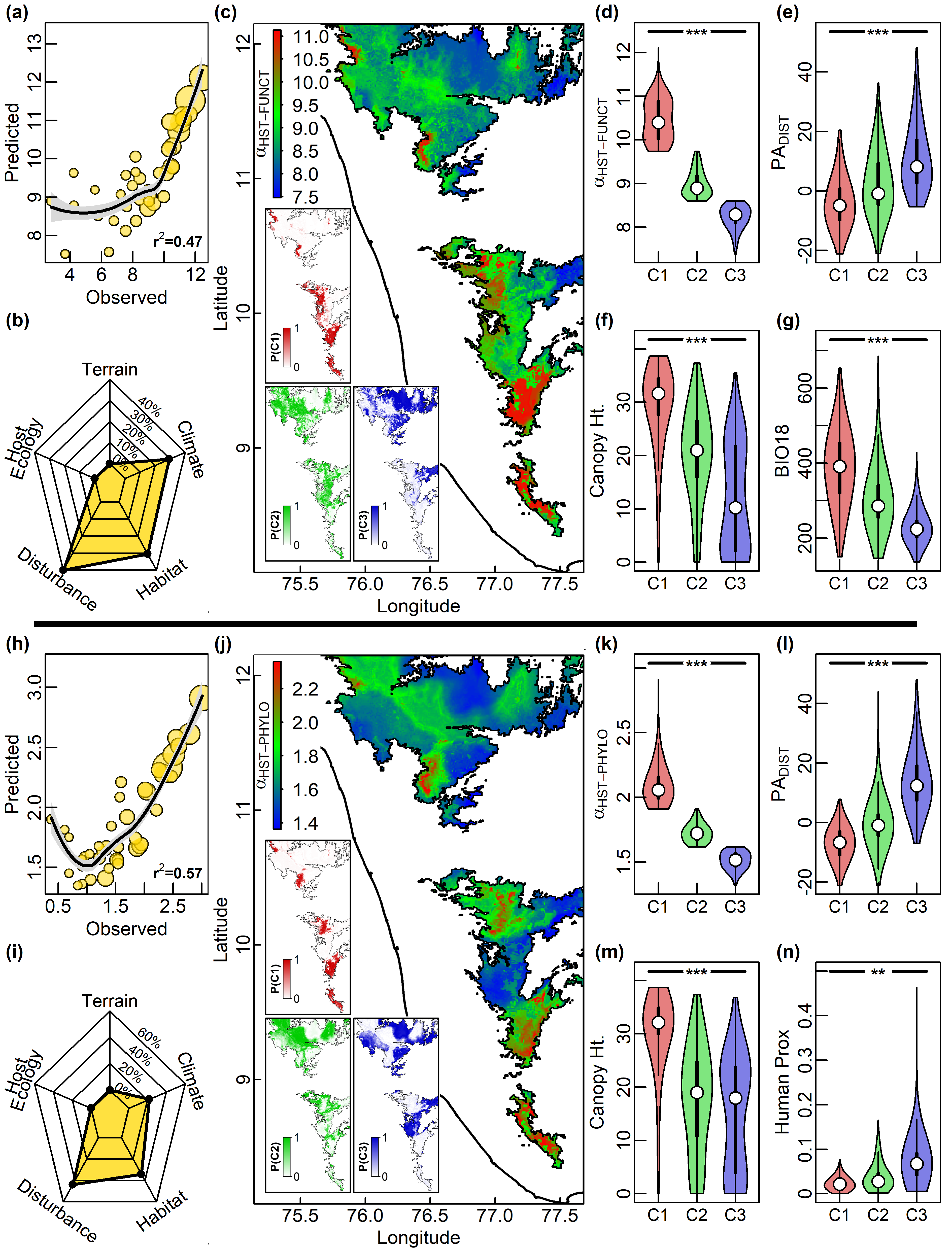

Spatial patterns of host functional α diversity (α_HST-FUNCT_) in the Western Ghats, Southern India, based on the best fit Random Forests Model (RFM) analysis (a-c). (a) Observed vs. predicted scatter plot for α_HST-FUNCT_, with size of points scaled relative to sample size in each site. (b) Relative importance of variables grouped into five major categories (terrain, climate- , habitat- , disturbance- and host ecology- related factors) in predicting α_HST-FUNCT_. (c) Predicted spatial patterns of α_HST-FUNCT_ across the Western Ghats study area. Inset maps show the geographical regions resulting from the Fuzzy C-Means (FCM) cluster analysis of α_HST-FUNCT_ showing the probability of membership in the identified clusters C1 (red), C2 (green) and C3 (blue), respectively. For each cluster (C1-C3), the distribution of values are plotted as violin plots (d-g) for: (d) The response variable (α_HST-FUNCT_), (e) distance to protected areas (PA_DIST_), (f) canopy height (Canopy-Ht), (g) precipitation of warmest quarter (BIO18). Spatial patterns of host phylogenetic α diversity (α_HST-PHYLO_) based on the best fit RFM analysis (h-j). (h) Observed vs. predicted scatter plot for α_HST-PHYLO_, with size of points scaled relative to sample size in each site. (i) Relative importance of variables grouped into five major categories (see Fig. 1d for details) in predicting α_HST-PHYLO_(j) Predicted spatial patterns of α_HST-PHYLO_ across the Western Ghats study area, with inset maps showing results of FCM analysis (as in Fig. 1e). For each cluster (C1-C3), the distribution of values are plotted as violin plots (k-m) for: (k) The response variable (α_HST-PHYLO_), (l) distance to protected areas (PA_DIST_), (m) canopy height (Canopy-Ht), (n) human proximity index (Human-Prox). For box plots, the thick horizontal line and box represent the median and quartiles, respectively. Whiskers extend to 1.5 times the inter-quartile range and open circles represent outliers. For violin plots, the white dot and thick line represent the median and quartiles, respectively. The thin lines extend to 1.5 times the inter-quartile range. Significant differences between groups in box and violin plots are are indicated by the following symbols: *** (*P* < 0.001), ** (*P* < 0.01), * (*P* < 0.05), ns (non-significant). See text for statistical test details.

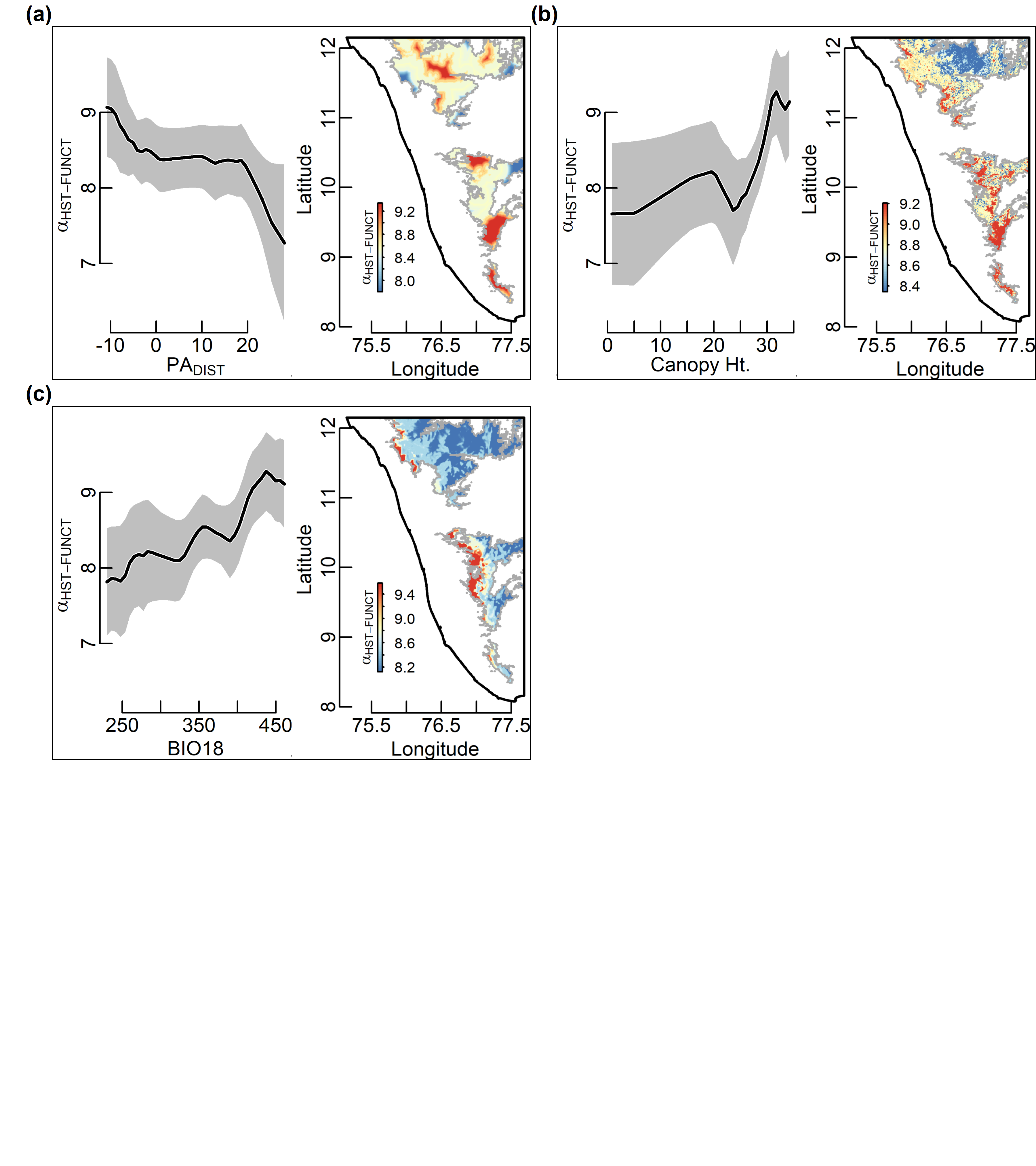

**Figure S06:** Partial dependence plots, based on the final random forest model for host functional α diversity (α_HST-FUNCT_) in the Western Ghats, Southern India. Separate figures are plotted for each independent variable in the random forest model, including: (a) distance to protected areas (PA_DIST_), (b) canopy height (Canopy-Ht), (c) precipitation of warmest quarter (BIO18). In each plot, the figure on the left shows the mean marginal influence of a particular independent variable on α_HST-FUNCT_ while holding the other independent variables constant. Alternatively, the figure on the right shows the spatially explicit partial predictions for α_HST-FUNCT_ while holding the other independent variables constant.

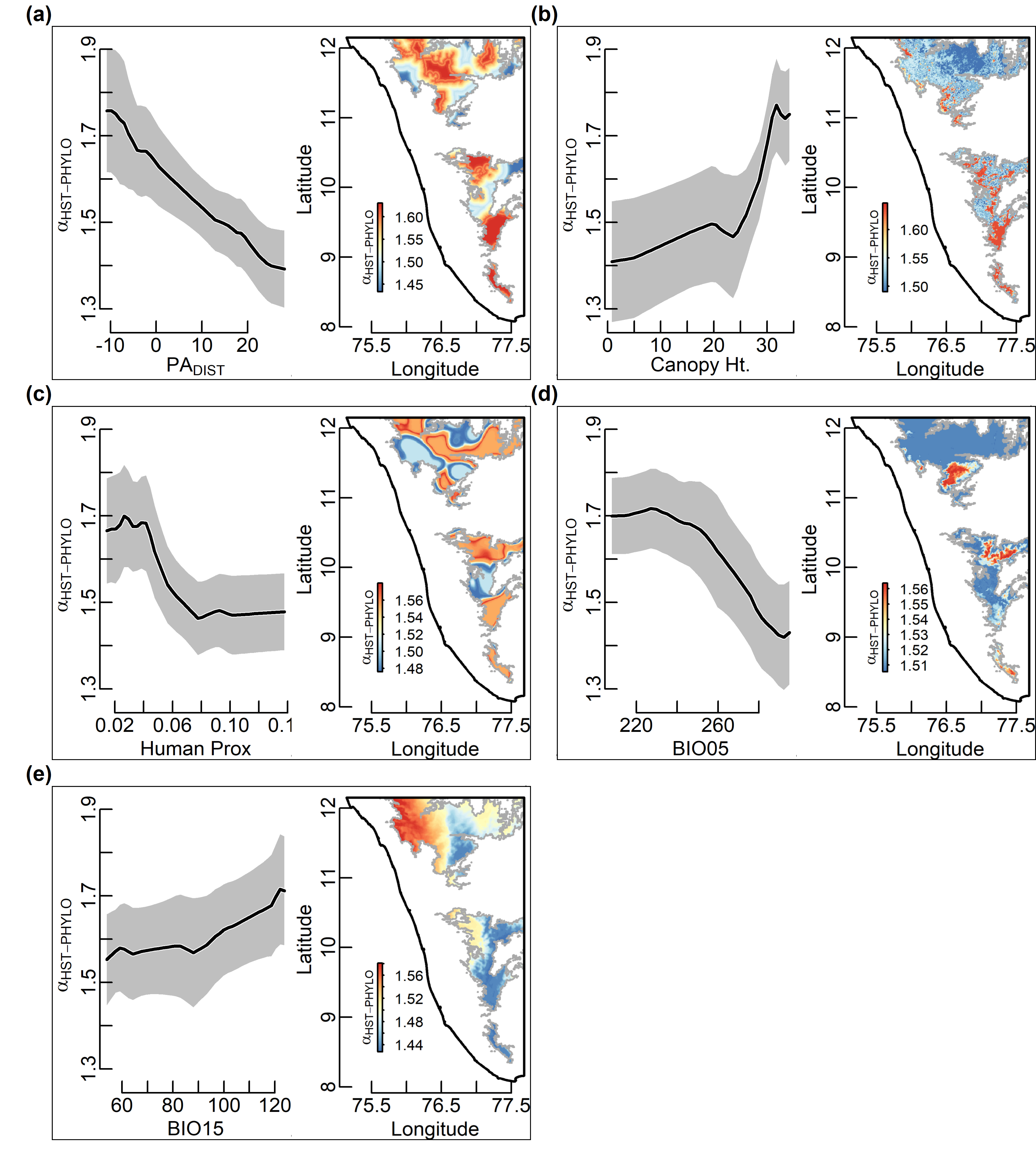

**Figure S07:** Partial dependence plots, based on the final random forest model for host phylogenetic α diversity (α_HST-PHYLO_) in the Western Ghats, Southern India. Separate figures are plotted for each independent variable in the random forest model, including: (a) distance to protected areas (PA_DIST_), (b) canopy height (Canopy-Ht), (c) human proximity index (Human-Prox), (d) max temperature of warmest month (BIO05), (e) precipitation seasonality (BIO15). In each plot, the figure on the left shows the mean marginal influence of a particular independent variable on α_HST-PHYLO_ while holding the other independent variables constant. Alternatively, the figure on the right shows the spatially explicit partial predictions for α_HST-PHYLO_ while holding the other independent variables constant.

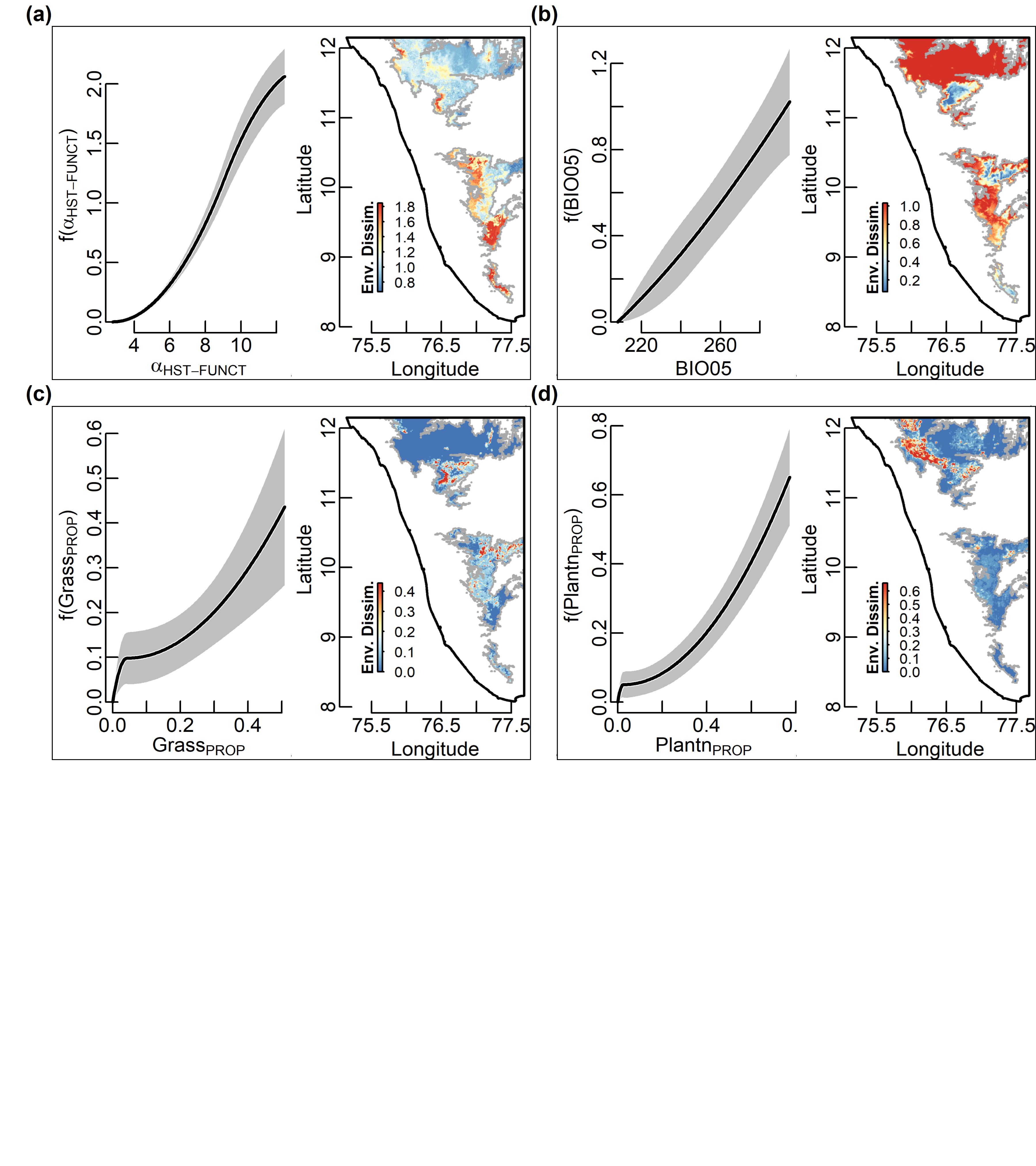

**Figure S08:** Partial dependence plots, based on the final generalized dissimilarity model (GDM) for *Haemoproteus* phylogenetic total β diversity (β_H-PHYLO_) in the Western Ghats, Southern India. Separate figures are plotted for each independent variable in the GDM, including: (a) host functional α diversity (α_HST-FUNCT_), (b) max temperature of warmest month (BIO05), (c) proportion grassland (Grass_PROP_), (d) proportion plantation (Plantn_PROP_). In each plot, the figure on the left shows the fitted splines from the GDM for β_H-PHYLO_ The height of the spline indicates the magnitude of diversity change along that gradient while holding all other variables constant (i.e. is a partial ecological distance). Alternatively, the figure on the right shows the spatially explicit partial ecological distance for each independent variable while holding the other independent variables constant.

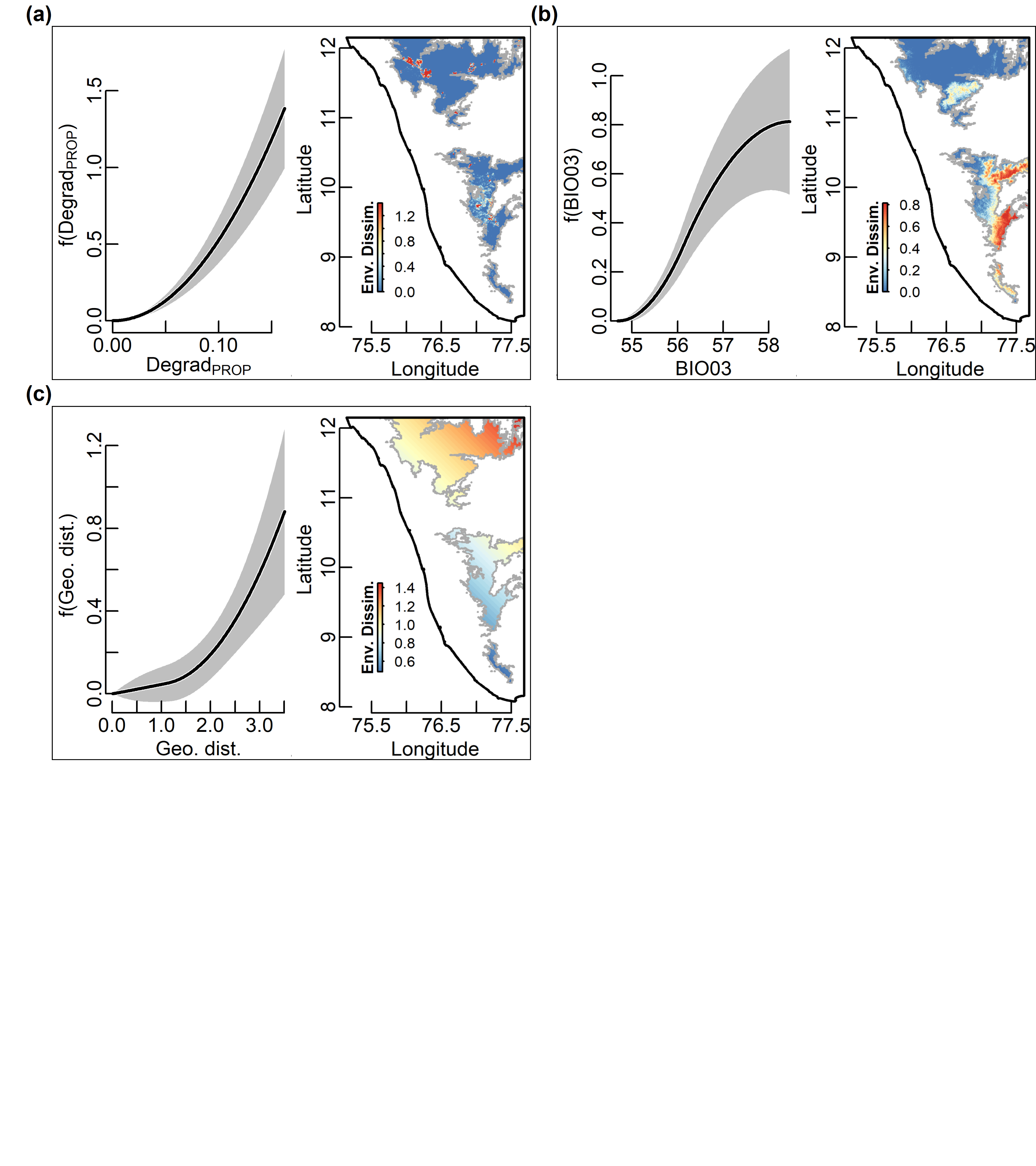

**Figure S09:** Partial dependence plots, based on the final generalized dissimilarity model (GDM) for *Plasmodium* phylogenetic total β diversity (β_P-PHYLO_) in the Western Ghats, Southern India. Separate figures are plotted for each independent variable in the GDM, including: (a) proportion degraded habitat (Degrad_PROP_), (b) isothermality (BIO03), (c) geographic distance (Geo-dist). In each plot, the figure on the left shows the fitted splines from the GDM for β_P-PHYLO_ The height of the spline indicates the magnitude of diversity change along that gradient while holding all other variables constant (i.e. is a partial ecological distance). Alternatively, the figure on the right shows the spatially explicit partial ecological distance for each independent variable while holding the other independent variables constant.

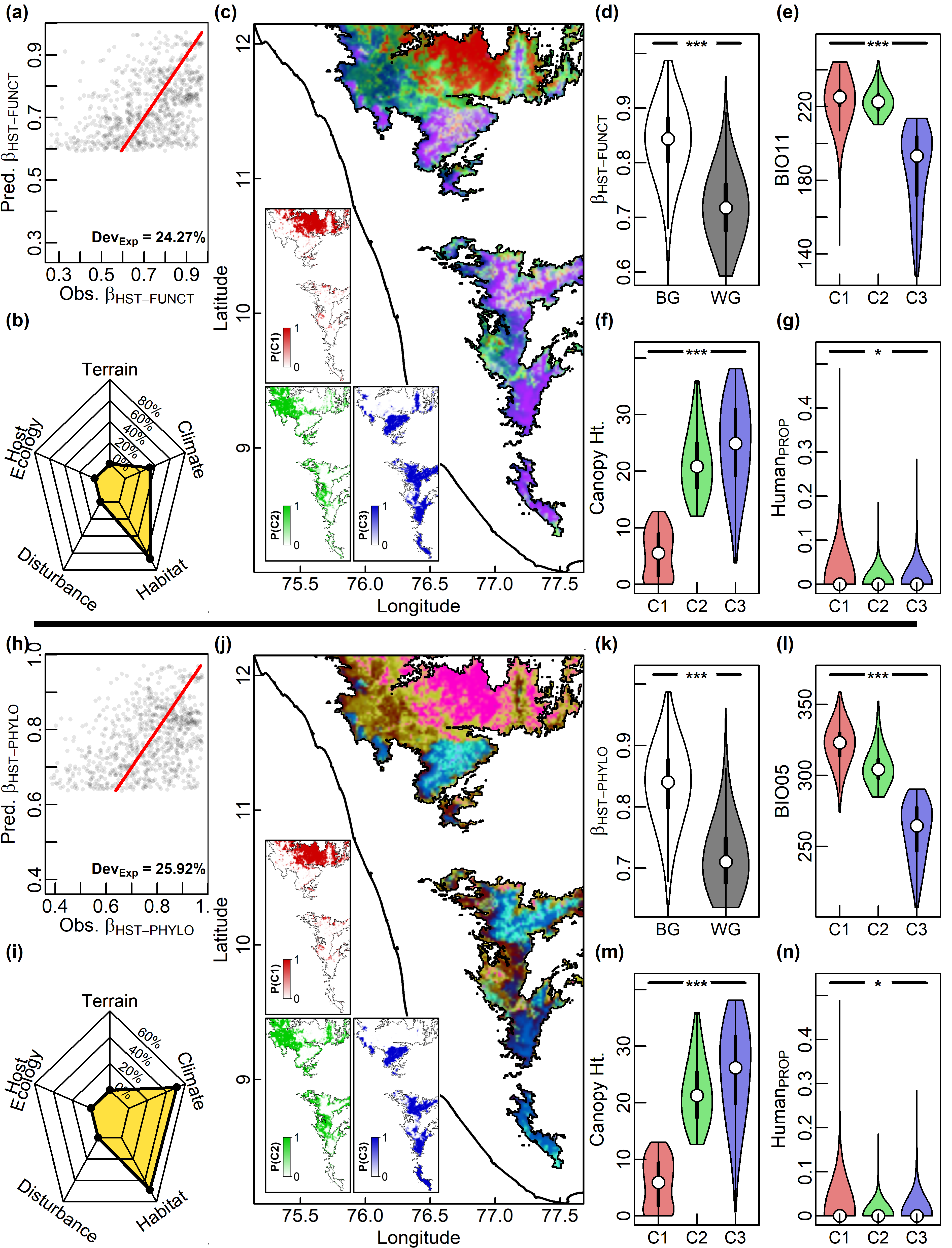

Spatial patterns of host functional β diversity (β_HST-FUNCT_) in the Western Ghats, Southern India based on the best fit generalized difference model (GDM) analysis (a-c). (a) Observed dissimilarity as a function of GDM-predicted dissimilarity for . Each site-pair is represented as a point and the line of is equality provided. (b) Relative importance of variables grouped into five major categories (terrain, climate- , habitat- , disturbance- and host ecology- related factors) in predicting . (c) Predicted spatial patterns of across the Western Ghats study area. Inset maps show the geographical regions resulting from the Fuzzy C-Means (FCM) cluster analysis of showing the probability of membership in the identified clusters C1 (red), C2 (green) and C3 (blue), respectively. For each cluster (C1-C3), the distribution of values are plotted as violin plots (d-g) for: (d) The response variable (β_HST-FUNCT_), (e) mean temperature of coldest quarter (BIO11), (f) canopy height (Canopy-Ht), (g) proportion settlements (Human_PROP_). Spatial patterns of host phylogenetic β diversity (β_HST-PHYLO_) based on the best fit generalized difference model (GDM) analysis (h-j). (h) Observed dissimilarity as a function of GDM-predicted dissimilarity for . Each site-pair is represented as a point and the line of is equality provided. (i) Variable importance of the predictors, including: (i) Relative importance of variables grouped into five major categories (terrain, climate- , habitat- , disturbance- and host ecology- related factors) in predicting . (j) Predicted spatial patterns of across the Western Ghats study area. Inset maps show the geographical regions resulting from the Fuzzy C-Means (FCM) cluster analysis of showing the probability of membership in the identified clusters C1 (red), C2 (green) and C3 (blue), respectively. For each cluster (C1-C3), the distribution of values are plotted as violin plots (k-n) for: (k) The response variable (β_HST-PHYLO_), (l) max temperature of warmest month (BIO05), (m) canopy height (Canopy-Ht), (n) proportion settlements (Human_PROP_). For box plots, the thick horizontal line and box represent the median and quartiles, respectively. Whiskers extend to 1.5 times the inter-quartile range and open circles represent outliers. For violin plots, the white dot and thick line represent the median and quartiles, respectively. The thin lines extend to 1.5 times the inter-quartile range. Significant differences between groups in box and violin plots are are indicated by the following symbols: *** (*P* < 0.001), ** (*P* < 0.01), * (*P* < 0.05), ns (non-significant). See text for statistical test details.

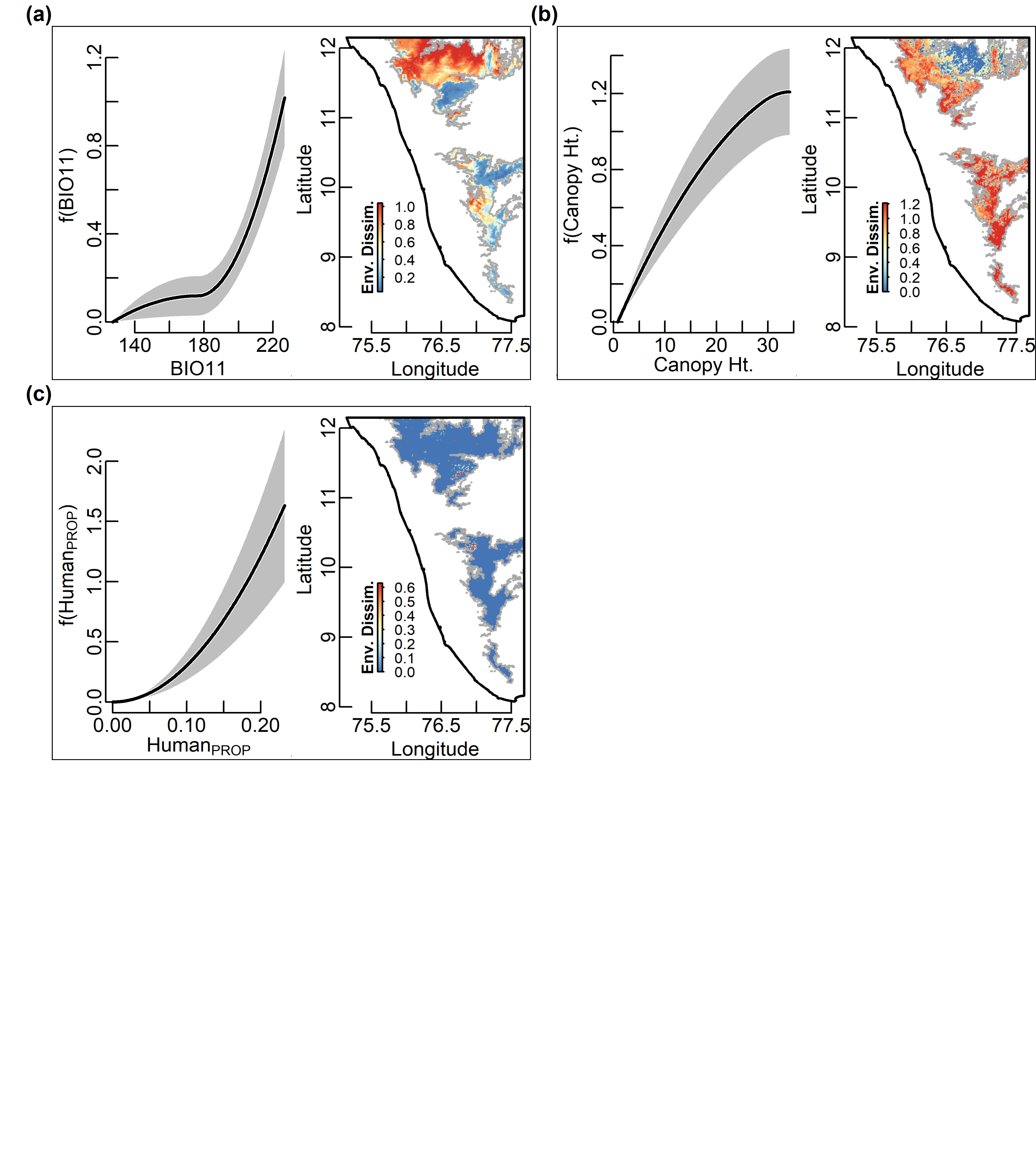

**Figure S11:** Partial dependence plots, based on the final generalized dissimilarity model (GDM) for host functional total β diversity (β_HST-FUNCT_) in the Western Ghats, Southern India. Separate figures are plotted for each independent variable in the GDM, including: (a) mean temperature of coldest quarter (BIO11), (b) canopy height (Canopy-Ht), (c) proportion settlements (Human_PROP_). In each plot, the figure on the left shows the fitted splines from the GDM for β_HST-FUNCT_ The height of the spline indicates the magnitude of diversity change along that gradient while holding all other variables constant (i.e. is a partial ecological distance). Alternatively, the figure on the right shows the spatially explicit partial ecological distance for each independent variable while holding the other independent variables constant.

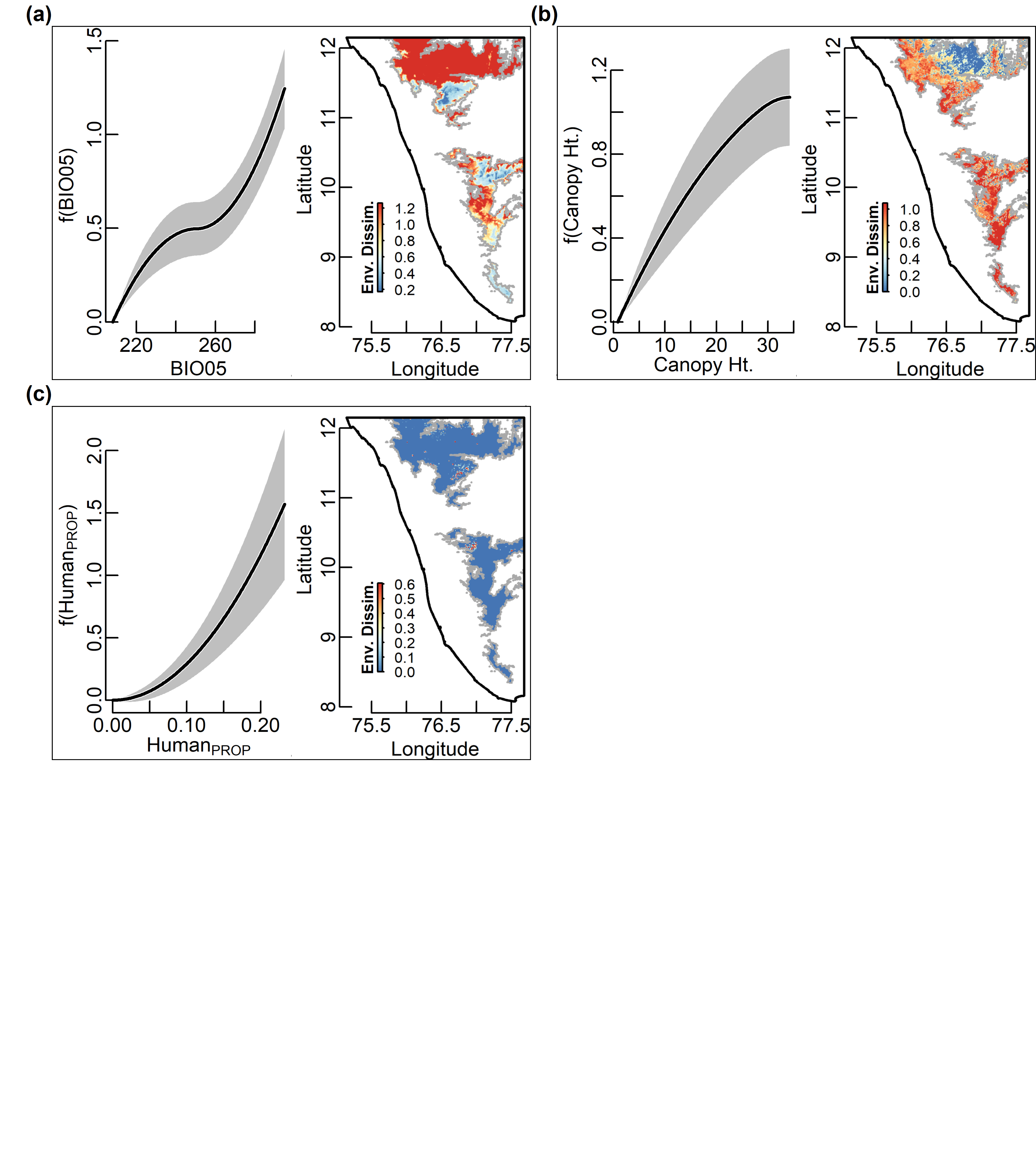

**Figure S12:** Partial dependence plots, based on the final generalized dissimilarity model (GDM) for host phylogenetic total β diversity (β_HST-PHYLO_) in the Western Ghats, Southern India. Separate figures are plotted for each independent variable in the GDM, including: (a) max temperature of warmest month (BIO05), (b) canopy height (Canopy-Ht), (c) proportion settlements (Human_PROP_). In each plot, the figure on the left shows the fitted splines from the GDM for β_HST-PHYLO_ The height of the spline indicates the magnitude of diversity change along that gradient while holding all other variables constant (i.e. is a partial ecological distance). Alternatively, the figure on the right shows the spatially explicit partial ecological distance for each independent variable while holding the other independent variables constant.

**SUPPLEMENTARY MATERIALS: TABLES**

**Table S1:** Sampling sites in each geographical region identified by a major and minor geographical location and elevation(m) at each trapping site for samples collected to study avian haemosporidians in the Shola sky island bird communities in Western Ghats (India). Each geographical region: I (Bababudan & Banasura hills), II (Nilgiri hills), III (Anamalai-Palni-Highwavies Hills), IV (Ashambu hills) corresponds to the major sky island group separated by three biogeographical barriers—Chaliyar Gap, Palghat Gap and Shencottah Gap (Fig. 1). The full data table can be found at: <https://doi.org/10.6084/m9.figshare.c.4509134.v2>

| **Region** | **Location** | **No. sites** | **Sample size** | **Latitude** | **Longitude** | **Elevation (m)** |
| --- | --- | --- | --- | --- | --- | --- |
| I | Vellarimala | 1 | 21 | 11.46 | 76.13 | 1732.23 |
| I | Chembra | 1 | 29 | 11.53 | 76.09 | 1622.50 |
| I | Banasura | 3 | 61 | 11.66 | 75.93 | 1590.98 |
| I | Ambalapara | 4 | 150 | 11.94 | 75.95 | 1418.99 |
| II | Sispara | 2 | 86 | 11.18 | 76.47 | 2105.28 |
| II | Silent valley | 9 | 147 | 11.20 | 76.55 | 1504.43 |
| II | Ooty | 1 | 15 | 11.39 | 76.69 | 2256.01 |
| III | Periyar | 3 | 25 | 9.56 | 77.15 | 874.89 |
| III | High Wavies | 3 | 141 | 9.58 | 77.33 | 1571.63 |
| III | Munnar | 3 | 189 | 10.13 | 77.09 | 2083.29 |
| III | Kodi | 6 | 99 | 10.23 | 77.43 | 2039.92 |
| III | Grasshills | 1 | 7 | 10.33 | 77.02 | 1566.09 |
| IV | Peppara | 5 | 202 | 8.68 | 77.19 | 1088.81 |
| Total (*or Mean*) |  | 42 | 1172 | *10.68* | *76.69* | *1650.39* |

**Table S2:** List of variables used for analyses

| **Variable** | **Description** |
| --- | --- |
| BIO01 | annual mean temperature |
| BIO02 | mean diurnal range |
| BIO03 | isothermality |
| BIO04 | temperature seasonality |
| BIO05 | max temperature of warmest month |
| BIO06 | min temperature of coldest month |
| BIO07 | temperature annual range |
| BIO08 | mean temperature of wettest quarter |
| BIO09 | mean temperature of driest quarter |
| BIO10 | mean temperature of warmest quarter |
| BIO11 | mean temperature of coldest quarter |
| BIO12 | annual precipitation |
| BIO13 | precipitation of wettest month |
| BIO14 | precipitation of driest month |
| BIO15 | precipitation seasonality |
| BIO16 | precipitation of wettest quarter |
| BIO17 | precipitation of driest quarter |
| BIO18 | precipitation of warmest quarter |
| BIO19 | precipitation of coldest quarter |
| PA_DIST_ | distance to protected areas |
| Human-Prox | human proximity index |
| Canopy-Ht | canopy height |
| Agricul_PROP_ | proportion cropland |
| Degrad_PROP_ | proportion degraded habitat |
| Forest_PROP_ | proportion forest |
| Grass_PROP_ | proportion grassland |
| Plantn_PROP_ | proportion plantation |
| Human_PROP_ | proportion settlements |
| Shrub_PROP_ | proportion shrub and savannah |
| Water_PROP_ | proportion water |
| Elevation | elevation |
| Flow-Acc | accumulation of water flow |
| Slope | steepness degree of inclination |
| Geo-dist | geographic distance |
| Geo-dist | geographic distance |
| α_HST-FUNCT_ | host functional α diversity |
| α_HST-PHYLO_ | host phylogenetic α diversity |
| α_HST-FUNCT_-or-α_HST-PHYLO_ | host α diversity |
| α_H-PHYLO_ | *Haemoproteus* phylogenetic α diversity |
| α_P-PHYLO_ | *Plasmodium* phylogenetic α diversity |
| α_H-PHYLO_-or-α_P-PHYLO_ | parasite α diversity |
| α_H-EVENNESS_ | *Haemoproteus* α eveness |
| α_P-EVENNESS_ | *Plasmodium* α eveness |
| β_HST-FUNCT_ | host functional total β diversity |
| β_HST-FUNCT-REPL_ | host functional replacement related β diversity |
| β_HST-FUNCT-RICH_ | host functional richness related β diversity |
| β_HST-PHYLO_ | host phylogenetic total β diversity |
| β_HST-PHYLO-REPL_ | host phylogenetic replacement related β diversity |
| β_HST-PHYLO-RICH_ | host phylogenetic richness related β diversity |
| β_H-PHYLO_ | *Haemoproteus* phylogenetic total β diversity |
| β_H-PHYLO-REPL_ | *Haemoproteus* phylogenetic replacement related β diversity |
| β_H-PHYLO-RICH_ | *Haemoproteus* phylogenetic richness related β diversity |
| β_P-PHYLO_ | *Plasmodium* phylogenetic total β diversity |
| β_P-PHYLO-REPL_ | *Plasmodium* phylogenetic replacement related β diversity |
| β_P-PHYLO-RICH_ | *Plasmodium* phylogenetic richness related β diversity |
| Pred-β_HST-FUNCT_ | predicted host functional total β diversity |
| Pred-β_HST-PHYLO_ | predicted host phylogenetic total β diversity |
| Pred-β_H-PHYLO_ | predicted *Haemoproteus* phylogenetic total β diversity |
| Pred-β_P-PHYLO_ | predicted *Plasmodium* phylogenetic total β diversity |
| Obs-β_HST-FUNCT_ | observed host functional total β diversity |
| Obs-β_HST-PHYLO_ | observed host phylogenetic total β diversity |
| Obs-β_H-PHYLO_ | observed *Haemoproteus* phylogenetic total β diversity |
| Obs-β_P-PHYLO_ | observed *Plasmodium* phylogenetic total β diversity |

**Table S3:** List of R packages used for analyses

| **Package** | **Version** | **Date** | **URL** |
| --- | --- | --- | --- |
| ape | 5.8-1 | Dec-10-2024 | <https://CRAN.R-project.org/package=ape> |
| arsenal | 3.6.3 | Jun-04-2021 | <https://CRAN.R-project.org/package=arsenal> |
| BAT | 2.9.6 | Feb-16-2024 | <https://CRAN.R-project.org/package=BAT> |
| caret | 6.0-94 | Mar-21-2023 | <https://CRAN.R-project.org/package=caret> |
| cluster | 2.1.6 | Nov-30-2023 | <https://CRAN.R-project.org/package=cluster> |
| corrplot | 0.92 | Nov-11-2021 | <https://CRAN.R-project.org/package=corrplot> |
| doParallel | 1.0.17 | Feb-07-2022 | <https://CRAN.R-project.org/package=doParallel> |
| foreach | 1.5.2 | Feb-02-2022 | <https://CRAN.R-project.org/package=foreach> |
| gdata | 3.0.1 | Oct-22-2024 | <https://CRAN.R-project.org/package=gdata> |
| gdm | 1.5.0-9.1 | Nov-17-2022 | <https://CRAN.R-project.org/package=gdm> |
| geosphere | 1.5-18 | Nov-13-2022 | <https://CRAN.R-project.org/package=geosphere> |
| ggplot2 | 4.0.2 | Feb-03-2026 | <https://CRAN.R-project.org/package=ggplot2> |
| iterators | 1.0.14 | Feb-05-2022 | <https://CRAN.R-project.org/package=iterators> |
| kableExtra | 1.4.0 | Jan-24-2024 | <https://CRAN.R-project.org/package=kableExtra> |
| knitr | 1.48 | Jul-07-2024 | <https://CRAN.R-project.org/package=knitr> |
| lattice | 0.22-6 | Mar-20-2024 | <https://CRAN.R-project.org/package=lattice> |
| Matrix | 1.7-3 | Mar-05-2025 | <https://CRAN.R-project.org/package=Matrix> |
| nlme | 3.1-164 | Nov-27-2023 | <https://CRAN.R-project.org/package=nlme> |
| pdp | 0.8.1 | Jun-07-2022 | <https://CRAN.R-project.org/package=pdp> |
| permute | 0.9-7 | Jan-27-2022 | <https://CRAN.R-project.org/package=permute> |
| picante | 1.8.2 | Jun-08-2020 | <https://CRAN.R-project.org/package=picante> |
| png | 0.1-8 | Nov-29-2022 | <https://CRAN.R-project.org/package=png> |
| ranger | 0.17.0 | Nov-08-2024 | <https://CRAN.R-project.org/package=ranger> |
| raster | 3.6-26 | Oct-12-2023 | <https://CRAN.R-project.org/package=raster> |
| RColorBrewer | 1.1-3 | Apr-03-2022 | <https://CRAN.R-project.org/package=RColorBrewer> |
| rgdal | 1.6-6 | Apr-18-2023 | <https://CRAN.R-project.org/package=rgdal> |
| rgeos | 0.6-4 | Jun-18-2023 | <https://CRAN.R-project.org/package=rgeos> |
| sf | 1.0-16 | Mar-24-2024 | <https://CRAN.R-project.org/package=sf> |
| sp | 2.2-0 | Feb-01-2025 | <https://CRAN.R-project.org/package=sp> |
| spatialreg | 1.3-4 | Jun-10-2024 | <https://CRAN.R-project.org/package=spatialreg> |
| spData | 2.3.1 | May-31-2024 | <https://CRAN.R-project.org/package=spData> |
| spdep | 1.3-5 | Jun-10-2024 | <https://CRAN.R-project.org/package=spdep> |
| stringr | 1.6.0 | Nov-04-2025 | <https://CRAN.R-project.org/package=stringr> |
| TeachingDemos | 2.13 | Feb-13-2024 | <https://CRAN.R-project.org/package=TeachingDemos> |
| tinytex | 0.52 | Jul-18-2024 | <https://CRAN.R-project.org/package=tinytex> |
| vegan | 2.6-6.1 | May-21-2024 | <https://CRAN.R-project.org/package=vegan> |

**Table S4:** Random forest model parameters and performance metrics

| **Dependent variable** | **Split rule** | **mTry** | **Min. node size** | **RMSE (**$\boldsymbol{\pm}$ **SD)** | **MAE (**$\boldsymbol{\pm}$ **SD)** | **R^2^ (**$\boldsymbol{\pm}$ **SD)** |
| --- | --- | --- | --- | --- | --- | --- |
| α_H-PHYLO_ | variance | 2 | 5 | 0.631 (0.313) | 0.526 (0.217) | 0.774 (0.307) |
| α_P-PHYLO_ | variance | 2 | 5 | 0.457 (0.140) | 0.398 (0.130) | 0.622 (0.376) |
| α_HST-FUNCT_ | extratrees | 3 | 5 | 2.276 (0.907) | 1.787 (0.733) | 0.474 (0.332) |
| α_HST-PHYLO_ | extratrees | 3 | 5 | 0.576 (0.194) | 0.493 (0.152) | 0.572 (0.309) |

**Table S5:** Variable importance measures for the best-fit random forest models

| **Dependent variable** | **Independent variable** | **Scaled variable importance** |
| --- | --- | --- |
| α_H-PHYLO_ | α_HST-PHYLO_ | 1.000 |
|  | α_HST-FUNCT_ | 0.849 |
|  | PA_DIST_ | 0.277 |
| α_P-PHYLO_ | α_HST-FUNCT_ | 0.973 |
|  | PA_DIST_ | 0.903 |
|  | BIO17 | 1.000 |
|  | BIO01 | 0.697 |
|  | Degrad_PROP_ | 0.469 |
|  | BIO04 | 0.589 |
| α_HST-FUNCT_ | PA_DIST_ | 1.000 |
|  | Canopy-Ht | 0.766 |
|  | BIO18 | 0.741 |
| α_HST-PHYLO_ | Canopy-Ht | 0.648 |
|  | PA_DIST_ | 1.000 |
|  | Human-Prox | 0.621 |
|  | BIO05 | 0.494 |
|  | BIO15 | 0.331 |

**Table S6:** Generalized dissimilarity best model assessment

| **Dependent variable** | **Model deviance** | **Deviance.explained (%)** | **Model P value** |
| --- | --- | --- | --- |
| β_HST-PHYLO_ | 22.326 | 25.919 | <0.001 |
| β_HST-FUNCT_ | 27.825 | 24.272 | <0.001 |
| β_H-PHYLO_ | 6.386 | 53.638 | <0.001 |
| β_P-PHYLO_ | 27.529 | 14.168 | <0.001 |

**Table S7:** Variable importance measures for the best-fit Generalized dissimilarity models

| **Dependent variable** | **Independent variable** | **Scaled variable importance** | **P value** |
| --- | --- | --- | --- |
| β_HST-PHYLO_ | BIO05 | 100.000 | 0.010 |
|  | Canopy-Ht | 47.214 | 0.040 |
|  | Human_PROP_ | 43.235 | 0.060 |
| β_HST-FUNCT_ | BIO11 | 100.000 | 0.010 |
|  | Canopy-Ht | 71.823 | 0.060 |
|  | Human_PROP_ | 55.512 | 0.070 |
| β_H-PHYLO_ | α_HST-FUNCT_ | 100.000 | <0.001 |
|  | BIO05 | 21.460 | <0.001 |
|  | Plantn_PROP_ | 9.724 | 0.040 |
|  | Grass_PROP_ | 9.339 | 0.100 |
| β_P-PHYLO_ | Degrad_PROP_ | 100.000 | 0.030 |
|  | BIO03 | 51.438 | 0.050 |
|  | Geographic | 45.015 | 0.040 |
